# Natural variation in Arabidopsis responses to *Plasmodiophora brassicae* reveals an essential role for RPB1

**DOI:** 10.1101/2022.10.06.511079

**Authors:** Juan Camilo Ochoa, Soham Mukhopadhyay, Tomasz Bieluszewski, Malgorzata Jedryczka, Robert Malinowski, William Truman

## Abstract

Despite the identification of clubroot resistance genes in various Brassica crops our understanding of the genetic basis of immunity to *Plasmodiophora brassicae* infection in the model plant *Arabidopsis thaliana* remains limited. To address this issue we performed a screen of 142 natural accessions and identified 11 clubroot resistant Arabidopsis lines. Genome wide association analysis identified several genetic loci significantly linked with resistance. Three genes from two of these loci were targeted for deletion by CRISPR/Cas9 mutation in resistant accessions Est-1 and Uod-1. Deletion of *Resistance to Plasmodiophora brassicae 1* (*RPB1*) rendered both lines susceptible to the *P. brassicae* pathotype P1+. Further analysis of *rpb1* knock-out Est-1 and Uod-1 lines showed that the RPB1 protein is required for activation of downstream defence responses, such as the expression of phytoalexin biosynthesis gene *CYP71A13*. RPB1 has no known functional domains or homology to previously characterised proteins. The clubroot susceptible Arabidopsis accession Col-0 lacks a functional *RPB1* gene; when Col-0 is transformed with *RPB1* expression driven by its native promoter it is capable of activating *RPB1* expression in response to infection but this is not sufficient to confer resistance. Constitutive over-expression of *RPB1* in Col-0 leads to drastically reduced growth and activation of stress-responsive genes. Furthermore, we found that transient expression of *RPB1* in *Nicotiana tabacum* induced programmed cell death in leaves. We conclude that RPB1 is a critical component of the defence response to *P. brassicae* infection in Arabidopsis, acting downstream of pathogen recognition but required for the elaboration of effective resistance.

## INTRODUCTION

Clubroot disease, caused by the soil-borne, obligate biotroph, protist *Plasmodiophora brassicae* (Woronin), is a significant challenge to the production of oilseed rape and other brassica crops. This intracellular pathogen colonises the roots and hypocotyls of infected plants, manipulating their growth and development patterns to drive the formation of large, unstructured galls (Malinowski et al., 2019). Resting spores of *P. brassicae* are highly robust and can remain viable in contaminated soil for up to 20 years with a half-life of 3.6 years (Wallenhammar, 1996). When stimulated to germinate, primary zoospores encyst on the root hairs and epidermis of the host where they form primary plasmodia and multiply before releasing secondary zoospores which go on to infect the main root cortex (Liu et al., 2020b). Formation of clubroot galls during this secondary infection is characterised by the prolonged maintenance of the mitotic state in cambial progeny cells leading to hyperplasia. Further disease progression is associated with hypertrophy of the pathogen colonised underground parts, facilitated by host endoreduplication and cell expansion processes (Olszak et al., 2019). In addition to disruption of cell-cycle control, the growth of clubroot galls is fuelled via the manipulation of vascular development patterns with a reduction of xylem tissue and increase in the number and complexity of phloem poles (Walerowski et al., 2018). While these changes increase the transfer of photoassimilates to the gall the impact on water uptake and transpiration causes wilting in the above-ground tissue which significantly impacts the yield of clubroot infected crops.

Management of clubroot disease in infected areas involves the use of crop rotation and sanitation of farm equipment; while soil amendment techniques such as liming are popular, the use of microbicidal chemical control is limited due to expense and environmental concerns (Peng et al., 2014). Recently, the use of biocontrol agents to suppress *P. brassicae* has garnered increased attention; however, the most commonly adopted approach is the use of resistant cultivars (Diederichsen et al., 2009; Struck et al., 2022). The majority of sources of clubroot resistance have been identified in Brassica species, particularly *B. rapa* and *B. oleracea* (Ramzi et al., 2018; Mehraj et al., 2020). Two clubroot resistance genes have been cloned so far, *CRa* and *Crr1a*; both are TIR-NBS-LLR (Toll/interleukin-1 receptor-like – nucleotide-binding site – leucine-rich repeat) domain containing genes identified in *B. rapa* (Ueno et al., 2012; Hatakeyama et al., 2013). The first clubroot resistant cultivar of oilseed rape to be commercially released was the ‘Mendel’ cultivar which was generated from a re-synthesis of *B. napus* using clubroot resistant *B. rapa* and *B. oleracea;* one of the sources of resistance was later shown to be the *CRa/CRb* repeat identified in *B. rapa* (Diederichsen et al., 2006; Fredua-Agyeman and Rahman, 2016). The distribution of *P. brassicae* pathotypes capable of overcoming the resistance conferred by cv. ‘Mendel’ has recently expanded in Germany, a situation also found in Canada with the emergence of increasing virulence against commonly used clubroot resistant varieties of oilseed rape (Strelkov et al., 2016; Zamani-Noor et al., 2022). In common with other biotrophic pathogens, the response to *P. brassicae* infection in clubroot resistant hosts is associated with salicylic acid mediated signalling pathways (Lemarié et al., 2015a; Galindo-González et al., 2020). The importance of this hormone is reinforced by the characterisation of the *P. brassicae* virulence factor *PbBSMT* which methylates host salicylic acid in order to suppress defence responses (Bulman et al., 2019).

The model plant Arabidopsis can be infected by *P. brassicae* and surveys of the available natural variation have identified clubroot resistant and tolerant accessions. Fuchs and Sacristan (1996) tested 30 ecotypes for their response to four different isolates and identified Tsu-0 and Ze-0 as resistant to isolate e. This resistance was found to be associated with a dominant allele mapped to chromosome 1, that they designated *Resistance to Plasmodiophora brassicae 1* (*RPB1*). Alix et al, (2007) observed partial clubroot resistance to isolates eH and Ms6 in the accession Bur-0; analysis of crosses between Col-0 and Bur-0 identified four additive QTLs accounting for 33.8% of the resistance coming from Bur-0 (Jubault et al., 2008). Subsequent investigation of these QTLs associated the partial resistance to *P. brassicae* with accumulation of the phytoalexin camalexin and tolerance of trehalose-induced toxicity (Gravot et al., 2011; Lemarié et al., 2015b). The interaction of 84 Arabidopsis accessions and Canadian *P. brassicae* pathotypes 2, 3, 5, and 6 identified clubroot resistance in accessions Ct-1, Pu2-23, Ws-2 and Sorbo, with accessions carrying strong resistance to no more than one of the pathotypes tested (Sharma et al., 2013). Despite the identification of clubroot resistance genes in various Brassica crops no such resistance genes have been characterised in Arabidopsis.

In this study, analysis of the interaction between a set of 142 natural inbred accessions of Arabidopsis and a P1+ pathotype of *P. brassicae* originating from Poland identified several resistant Arabidopsis lines. Candidate genes selected from the GWAS analysis of this phenotype data were validated through the generation of knock-out mutations by CRISPR/Cas9 genome editing. Of the candidate genes evaluated, *RPB1* was found to be necessary for clubroot resistance though transformation of susceptible accession Col-0 revealed that it was not sufficient to confer full resistance.

## RESULTS

### Eleven clubroot resistant lines were identified in a screen of 142 natural Arabidopsis accessions

In order to survey the interaction of *P. brassicae* and different Arabidopsis natural accessions, we first optimised the inoculation and quantification of clubroot disease progress in the reference accession Col-0. 17-day old plants were inoculated with a high dose of *P. brassicae* spores (2 x 10^6^) using a P1+ pathotype (Some et al., 1996; Zamani-Noor et al., 2022) collected from infested oilseed rape in Poland. Quantitative PCR, targeting the *P. brassicae* gene *Pb18S*, was used to determine pathogen DNA levels in the hypocotyls and upper roots of inoculated plants. The earliest stage when infection could be reliably detected was 10 days post inoculation (dpi) (Supp. Fig. 1B). At this time the symptoms of infection were barely detectable from observation of the roots (Supp. Fig. 1A). Pathogen DNA amounts relative to host DNA increased exponentially until 16 dpi and then plateaued while the expansion of galls continued up to 28 dpi. The 19 dpi time-point was selected for assaying the collection of Arabidopsis accessions as a time-point during infection that *P. brassicae* DNA could be reproducibly quantified in various accessions. In addition to the quantification of relative pathogen DNA levels, the width of the galls were measured at the thickest point, and the disease index (DI) was calculated from a visual classification of clubroot symptom development (Ludwig-Müller et al., 2017). A collection of 142 accessions comprising 118 accessions sequenced in the 1001 genomes project (Alonso-Blanco et al., 2016) and 111 accessions genotyped as part of the RegMap panel (Horton et al., 2012) were tested with 5 separate inoculations. For the majority of accessions at least 21 plants were inoculated and scored, qPCR quantification was performed on DNA extracted from pools of three galls. Some accessions were affected by poor germination and were only included in a subset of inoculations, the minimum number of plants evaluated was 6 (Supp. Table 1). There was a strong degree of correlation between the DI score and the width of the gall (Spearman Rho = 0.81), the qPCR quantification was less strongly correlated with Spearman Rho scores of 0.57 and 0.52 respectively (Figure 1A). The majority of the accessions exhibited some degree of susceptibility to the P1+ pathotype used, only 11 accessions could be characterised as clearly resistant to infection (Figure 1A). Despite the lack of gall development in these accessions *P. brassicae* DNA could be amplified from all samples. Two outliers appear in the dataset, Pro-0 and Var-2-6. Pro-0, despite accumulating the highest levels of *P. brassicae* DNA relative to host DNA, developed galls that were significantly smaller than Col-0 and Kas-1 and lower in DI score (86.5 compared to 96.7 for Col-0 and 97.5 for Kas-1) (Supp. Fig. 2A). Var-2-6 did not exhibit any gall expansion despite strong symptoms of infection and a reasonably high pathogen DNA titre, the growth of this accession was severely affected by infection (Supp. Fig. 2B). The accessions determined to be resistant to clubroot disease (with a *Pb18S / AtSK11* log_2_ ratio less than 6 and DI score less than 55) include Fab-4, Est-1, Uod-1, Durh-1, HR-10, Mrk-0, Tsu-1, Bil-7, N13, along with Pu2-23, which was previously shown to be resistant to a Canadian pathotype, and also Tsu-0, previously observed to be resistant to *P. brassicae* isolate e (Fuchs and Sacristan 1996; Sharma et al., 2013). Within the resistant accessions significant differences in pathogen DNA accumulation could be discriminated with Fab-4 being the most restrictive followed by Est-1 and Uod-1 (Figure 1A, Supp. Table 1).

**Figure 1.**
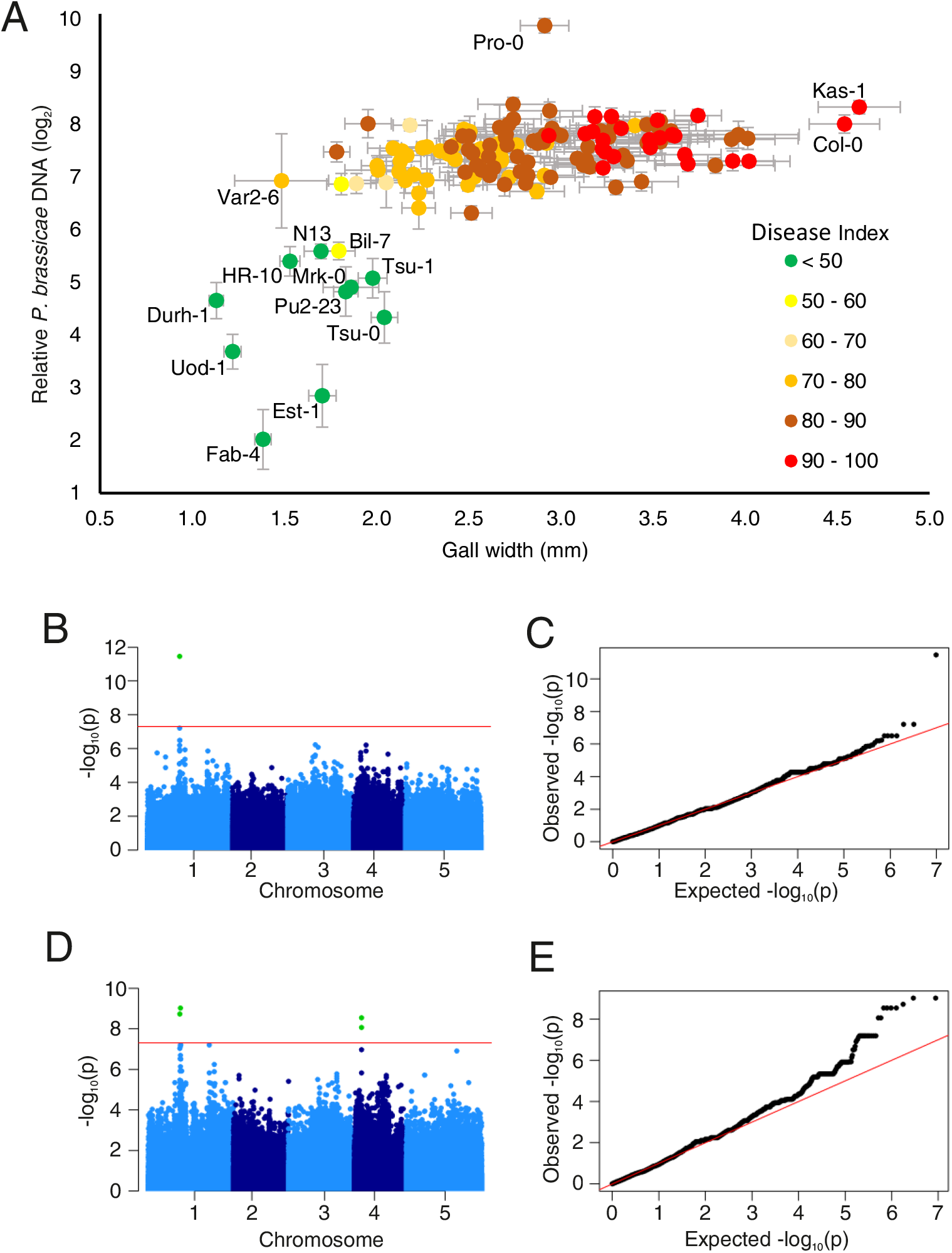
Resistance to *P. brassicae* is associated with three loci in the Arabidopsis genome. A) Clubroot phenotype scoring for 142 Arabidopsis accessions, relative pathogen DNA titre (*Pb18S/AtSK11*) is plotted against maximum gall width and coloured by DI score. For DNA quantification 3 plants were pooled for extraction 4 ≤ n ≤ 10, for width 12 ≤ n ≤ 30. Error bars indicate the standard error. B & C) GWA-Portal AMM analysis of pathogen DNA quantification for imputed genotypes for 140 accessions. D & E) Manhattan plot and Q-Q plot for easyGWAS EMMAX analysis for DI scores of 1001 genomes data for 118 accession. For B & D the red line indicates the threshold for significance a Bonferroni adjusted p-value < 0.05.

### Genome wide association analysis links clubroot resistance with two loci on chromosome 1

Genome wide association (GWA) analysis was applied to the dataset using two online toolsets for interrogating natural variation in Arabidopsis: easyGWAS (https://easygwas.ethz.ch/) (Grimm et al., 2017) and GWA-Portal (https://gwas.gmi.oeaw.ac.at) (Seren et al., 2013). The subsets of accessions covered by the 1001 genomes sequencing data and the RegMap genotyping data were analysed separately using both the GWA-Portal AMM and easyGWAS EMMAX algorithms; with GWA-Portal the imputed set of SNPs covering 141 accessions from both pools was also used. For both the gall width and relative pathogen DNA quantification phenotype data only one locus was found to be significantly associated with resistance at position 11515504 on Chromosome 1, whether the imputed genotype data or the RegMap data were used (Figure 1B, Table 1). In the analysis of the Disease Index phenotype data this SNP and two closely adjacent SNPs were identified within the imputed genotype dataset along with SNPs at an additional loci on chromosome 4 centred on position 2594401 with five neighbouring SNPs (Table 1). When the Disease Index phenotype was analysed in combination with the 1001 genomes sequence data these two loci from chromosome 1 and 4 were again highlighted along with an additional locus at position 11289740 on chromosome 1 (Figure 1D, Table 1). Pairing the Disease Index phenotype analysis with the RegMap genotyping two significant SNPS were identified, the frequently observed Chromosome 1 11515504 position and an additional SNP on Chromosome 3 at position 10506508 (Table 1). Of the four loci, the SNP with the highest significance score (Chromosome 1: 11515504) lies in the gene *AT1G32030* which falls within the region mapped to the *RPB1* locus previously associated with clubroot disease resistance (Fuchs and Sacristan, 1996; Arbeiter et al., 2002). The second SNP on chromosome 1 lies in the LRR domain of the TIR-NBS-LRR resistance gene *Resistance to Albugo candida 1 (RAC1) (AT1G31540)*; an allele of *RAC1* from the accession Ksk-1 has been shown to confer resistance to the oomycete pathogen *Albugo candida* (Borhan et al., 2004). The locus with multiple significant SNPs on chromosome 4 lies in the promoter region upstream of the genes *Wound-Induced Polypeptide 2* (*WIP2*) (*AT4G05070*) and *AT4G05071*. Sequencing of this region in the resistant accessions Uod-1, HR-10, Pu2-23 and Tsu-0 confirmed the presence of multiple SNPs, a 4 bp insertion and 9 bp deletion in the resistant accessions in addition to a larger rearrangement where 291 bp of the Col-0 sequence was absent in the resistant accessions, which instead contained 93 bp sequence that was poorly aligned with the Col-0 reference (Supp. Fig. 3A). No other potential open reading frames were observed as a consequence of these polymorphisms relative to Col-0. Expression of *AT4G05071* and *WIP2* was detected 7 dpi but no response to *P. brassicae* infection was observed, expression levels were higher in the susceptible Col-0 accession than the resistant Est-1 for *WIP2* (Supp. Fig. 3B, C). The significant SNP on Chromosome 3 is in the first exon of the cellulose synthase like gene *CSLC04* (*AT3G28180*). Based on the previous literature linking the SNP in *AT1G32030* with resistance to clubroot disease and the role of *RAC1* in mediating disease resistance these two loci were selected for further functional characterisation, leaving the region upstream of *WIP2* and the cellulose synthase gene for future investigation.

**Table 1.** SNPs significantly associated with different clubroot phenotype data (-log_10_ p-value).

| Algorithm | GWA-Portal AMM |  |  | easyGWAS EMMAX |  |
| --- | --- | --- | --- | --- | --- |
| Genotype data | Imputed | 1001 | 250K | 1001 | 250K |
| Gall width | Chr1: 11515504<br>(8.85) | - | Chr1: 11515504<br>(8.47) | - | Chr1: 11515504<br>(8.96) |
| Relative *P. brassicae*<br>DNA quantification | Chr1: 11515504<br>(11.46) | - | Chr1: 11515504<br>(11.86) | - | Chr1: 11515504<br>(11.67) |
| Disease Index | Chr1: 11515504<br>(13.13) | Chr1: 11289740<br>(8.07) | Chr1: 11515504<br>(12.36) | Chr1: 11513092<br>(9.02) | Chr1: 11515504<br>(12.74) |
|  | Chr1: 11513098<br>(10.26) | Chr4: 2594573<br>(8.79) | Chr3: 10506508<br>(7.41) | Chr1: 11513098<br>(9.02) | Chr3: 10506508<br>(7.54) |
|  | Chr1: 11512894<br>(8.38) | Chr4: 2594449<br>(8.79) |  | Chr1: 11289740<br>(8.73) |  |
|  | Chr4: 2594573<br>(9.51) | Chr4: 2594448<br>(8.79) |  | Chr4: 2594573<br>(8.54) |  |
|  | Chr4: 2594449<br>(9.51) | Chr4: 2594401<br>(8.79) |  | Chr4: 2594449<br>(8.54) |  |
|  | Chr4: 2594448<br>(9.51) | Chr4: 2594357<br>(8.16) |  | Chr4: 2594448<br>(8.54) |  |
|  | Chr4: 2594401<br>(9.51) | Chr4: 2594378<br>(8.16) |  | Chr4: 2594401<br>(8.54) |  |
|  | Chr4: 2594357<br>(8.87) |  |  | Chr4: 2594357<br>(8.06) |  |
|  | Chr4: 2594378<br>(8.87) |  |  | Chr4: 2594378<br>(8.06) |  |

### Deletion of *RAC1* does not affect clubroot resistance in Est-1 or Uod-1

Sequencing of *RAC1* in Uod-1, one of the resistant accessions carrying the C allele for chr1 position 11289740, confirmed the polymorphism, which gives rise to an alanine to serine substitution in the C-terminal region of the protein (Supp. Fig. 4A). The TIR domain encoded in the first exon, which is common to the various predicted protein isoforms, is more highly conserved than the LRR domains and the C-terminal region, which have previously been reported to be a hotspot for meiotic recombination (Serra et al., 2018). Sequencing of additional resistant accessions across the region flanking the SNP revealed extensive differences in the protein sequence compared to Col-0, with some variations shared by the resistant accessions Uod-1, HR-10 and Pu2-23, and some variants distinct to resistant accessions Est-1 and Tsu-0; susceptible accessions for which sequence information was available also displayed polymorphisms relative to the Col-0 sequence in this region (Supp. Fig. 4A). In order to generate null mutations of *RAC1* in the resistant accessions Est-1 and Uod-1, a CRISPR/Cas9 gene editing strategy was devised; these lines were transformed with a construct carrying two guide RNAs targeting the first exon of *RAC1*, spaced 105 bp apart. Mutations in *RAC1* were identified in plants from the T2 generation which no longer carried the CRISPR/Cas9 construct; sequencing of the targeted region revealed deletions ranging in size from 22 bp to 188 bp combined with other mutations (Supp. Table 2). Six homozygous mutants from the T3 generation, three in Est-1 and three in Uod-1 were selected for phenotyping; in each mutant the effect of the deletion introduced would be to truncate translation of any putative protein within the first 101 aa of the full length 1161 aa protein of RAC1 isoform AT1G31540.2. All six *rac1* mutant lines exhibited full resistance to *P. brassicae* infection, there was no difference in gall development with respect to DI score or hypocotyl expansion compared to wild-type Uod-1 and Est-1 and no significant difference in qPCR quantification of *P. brassicae* relative DNA levels compared to wild-type controls (Supp. Fig. 5A, B, C).

### Resistant accessions carry a conserved *RPB1* gene absent in Col-0

In 1996 the genetic locus *RPB1* conferring resistance to *P. brassicae* isolate e was identified in the accession Tsu-0 and linked to the *dis1* marker on chromosome 1 (Fuchs and Sacristan, 1996). Arbeiter et al. further refined the position of *RPB1* to a region of approximately 71 kb through analysis of a Tsu-0 x Cvi-0 mapping population in 2002 (Arbeiter et al., 2002). In 2009 Rehn and Siemens deposited two nucleotide sequences in the GenBank database annotated with *RPB1* and *RPB1-like* genes from the Tsu-0 and RLD-1 accessions designated FJ807885 and FN400762 respectively (unpublished). The sequence for accession Tsu-0 is annotated with two genes, *RPB1a* and *RPB1b*, both consisting of 447 bp single exon open reading frames, an additional pseudogene *RPB1-like-1* is also defined (Figure 2A). In the GenBank entry for accession RLD-1 a single copy of *RPB1*, also 447 bp in length, is defined in addition to orthologues for the Col-0 reference genes *AT1G32020* and *AT1G32030* and novel genes *RPB1-like-2, RPB1-like-3* and *RPB1-like-4*. The SNPs in *AT1G32030* that were found to be significantly associated with resistance to *P. brassicae* in this study are positioned within 1500 bp of the *RPB1* gene annotated in the sequence for RLD-1. Using primers derived from the sequences deposited to Genbank, which are significantly structurally different from the reference genome Col-0, overlapping fragments could be amplified from Est-1 and sequenced; a 10322 bp sequence for this region was assembled and found to match the structure of RLD-1 with only one ORF corresponding to *RPB1* as well as *RPB1-like-2, RPB1-like-3* and *RPB1-like-4* sequences (Figure 2A). Amplification and sequencing of portions of this region for Uod-1 revealed the same gene structure as Est-1 and RLD-1. The sequences of the *RPB1* genes from the *P. brassicae* resistant accessions in this study and those deposited to Genbank previously are highly conserved with 100% identity among nucleotide sequences for Est1 *RPB1*, Uod-1 *RPB1*, RLD-1 *RPB1*, Tsu-0 *RPB1a* and 99.3% identity with Tsu-0 *RPB1b*, for the amino acid sequences there is 100% identity (Supp. Fig. 6). The susceptible accession Col-0 contains a pseudogene *AT1G32049* with 54.2% identity to the other RPB1 nucleotide sequences. However, *RPB1* genes have been annotated in the genome assemblies of accessions that were found to be susceptible to *P. brassicae* in this study: An-1, Cvi-0, C24 and Kyoto all contain *RPB1* genes encoding putative proteins with 100% amino acid identity to those predicted for the resistant alleles with the exception of Cvi-0 which differs in one amino acid residue (Figure 2A, Suppl. Fig. 6) (Jiao and Schneeberger, 2020). The expression of *RPB1* and the *RPB1-like* genes was examined in Est-1 at 7 dpi, a time-point early in secondary infection previously shown to be critical for the restriction of *P. brassicae* growth in resistant cultivars (Liu et al., 2020a), only *RPB1* and *RPB1-like-4* were found to be expressed, no transcripts for *RPB1-like-2* or *RPB1-like-3* could be amplified. Both *RPB1* and *RPB1-like-4* were found to be significantly up-regulated in response to *P. brassicae* infection 7 dpi (Figure 2B).

**Figure 2.**
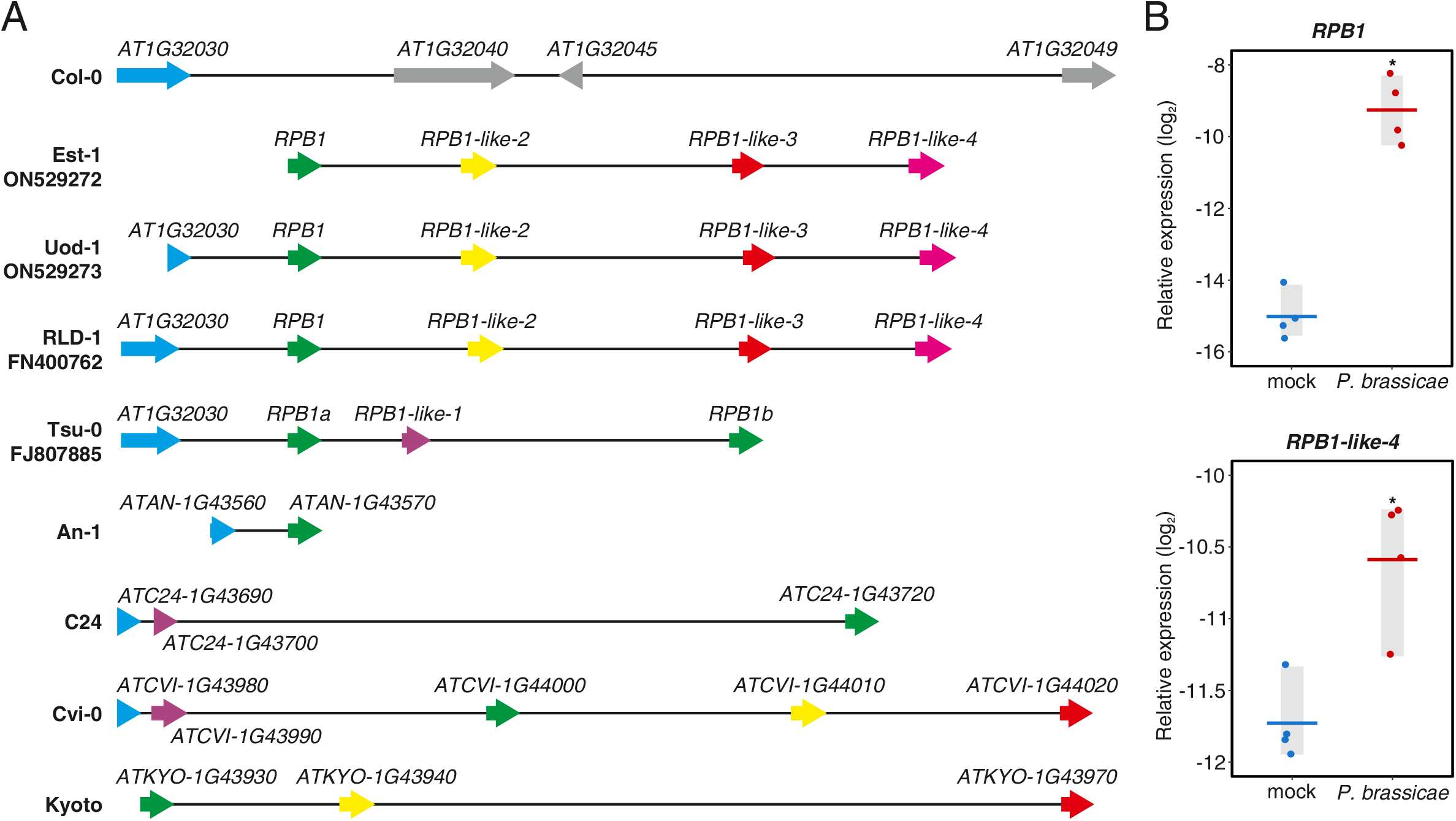
*RPB1* and *RPB1-like-4* genes respond to *P. brassicae* infection. A) Comparison of gene structures from annotated Arabidopsis genomes, Genbank sequences and Sanger sequencing of accessions Est-1 and Uod-1. Col-0, An-1, C24, Cvi-0 and Kyoto are susceptible to the P1+ pathotype tested while Est-1, Uod-1 and Tsu-0 are resistant, RLD-1 was not tested. Shared colours for genes indicate orthologous relationships identified by OrthoFinder analysis, grey indicates non-translated transposons and pseudogenes. B) Expression of *RPB1* and *RPB1-like-4* in Est-1 7 d after infection with *P. brassicae*. Expression is relative to *AT1G76030* and *AT3G48140*. Asterisks indicate significant differences in expression in response to treatment (p < 0.05). Each point represents one biological replicate (n = 4) of 10 plants, horizontal bars indicate the mean.

### *RPB1* is indispensable for resistance to resistance to *P. brassicae* in selected genotypes

In order to establish the potential function of *RPB1* and *RPB1-like-4* in mediating resistance to clubroot disease, both these genes were targeted for deletion using a CRISPR/Cas9 mutation approach. Est-1 and Uod-1 plants were transformed with constructs carrying two guide RNAs targeted to flank 352 bp and 316 bp regions within the single exon ORFs of *RPB1* and *RPB1-like-4* respectively. Additionally, a construct carrying all 4 guide RNAs was generated in order to target both genes simultaneously. Homozygous knock-out lines were generated in both accessions for *RPB1* and *RPB1-like-4* with deletion sizes ranging between 10 bp and 370 bp along with various insertions and substitutions (Table 2). For each gene in each accession two mutant alleles predicted to encode missense proteins with early truncation could be identified. For the dual targeting of *RPB1* and *RPB1-like-4* one homozygous double mutant was identified in the Uod-1 background with a 372 bp deletion in *RPB1* and single nucleotide deletions and insertions in *RPB1-like-4*, these are predicted to result in missense expression from aa 16 and aa 5, resulting in premature termination of translation from aa 24 and aa 11, respectively (Table 2). All nine mutant lines were tested for susceptibility to clubroot disease with quantitative evaluation of *P. brassicae* growth 19 dpi. The four lines with single mutations in *RPB1* were clearly susceptible to *P. brassicae* infection (Figure 3A), with significantly higher accumulation of pathogen DNA and expansion in gall width compared with wildtype controls (Figure 3B, C). These metrics for infection were not significantly different from the susceptible Col-0 controls, while the disease index scores ranged from 86.5 to 97% (Figure 3D). For the single mutations in *RPB1-like-4* no increased susceptibility was observed, in terms of relative *P. brassicae* DNA levels. While the gall expansion in Uod-1 mutant *rpb1-like-4-1* was slightly greater than wild-type Uod-1, the other three mutants were indistinguishable from the wildtype controls. Disease index scores for the mutants remained in the range 26.9 to 44.2%.

**Figure 3.**
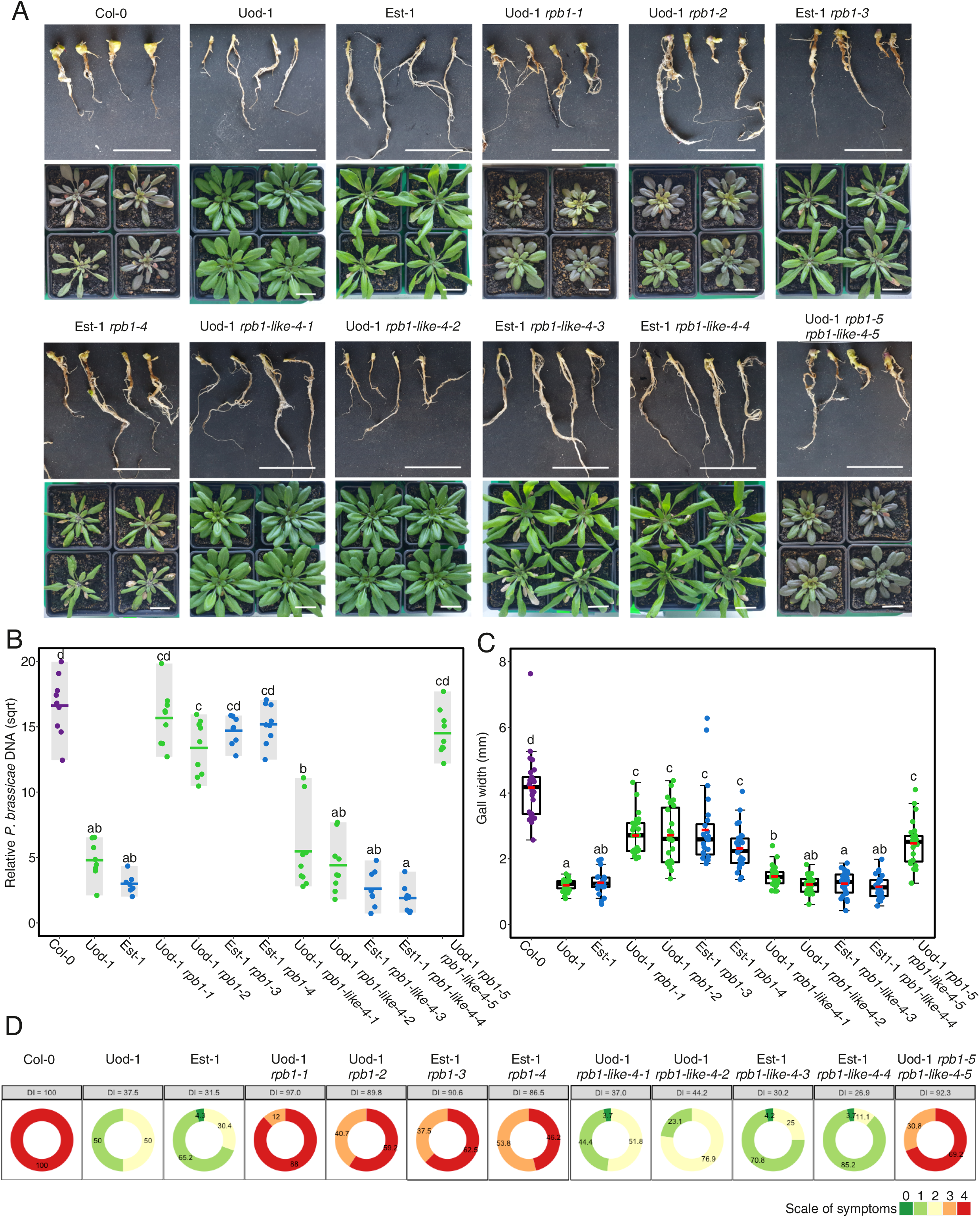
Deletion of *RPB1* compromises clubroot resistance in Est-1 and Uod-1 accessions. A) Gall and rosette symptoms of *P. brassicae* infection 19 dpi for *rpb1, rpb1-like-4* and *rpb1 rpb1-like-4* knock-out mutants. White bars = 2 cm. B) Relative pathogen DNA titre (*Pb18S/AtSK11*) 19 dpi, points indicate biological replicates of 3 plants, horizontal lines indicate the means, different letters indicate statistically significant differences (Tukey, BH adjusted p < 0.05), 7 ≤ n ≤ 9. C) Gall width measurements for individual plants pooled for B, red lines indicate the mean, black boxes the lower quartile, median and upper quartile, whiskers the 1.5 IQR, 21 ≤ n ≤ 27, different letters indicate statistically significant differences (Games-Howell, BH adjusted p < 0.05). D) DI score for the symptoms of galls in B & C, the percentage of plants assigned to each symptom class is shown on each segment.

**Table 2.** Mutations generated in *RPB1* and *RPB1-like-4*.

| Name | Background | Impact on *RPB1* gene (single exon of 444 bp) | Impact on RPB1 protein (148 aa) |
| --- | --- | --- | --- |
| *rpb1-1* | Uod-1 | c.47_418del | p.(Gly16Alafs*25) |
| *rpb1-2* | Uod-1 | c.47_419del | p.(Gly16Alafs*?) |
| *rpb1-3* | Est-1 | c.44_55del, c.420_421insC | p.(Gly16Glufs*53) |
| *rpb1-4* | Est-1 | c.46_47insT, c.385_425del | p.(Gly16Valfs*17) |
| Name | Background | Impact on *RPB1-like-4* gene (single exon of 477 bp) | Impact on RPB1-like-4 protein (159 aa) |
| *rpb1-like4-1* | Uod-1 | c.14_121del | p.(Gly5Valfs*124) |
| *rpb1-like4-2* | Uod-1 | c.14_17del, c.327_364del | p.(Gly5Valfs*11) |
| *rpb1-like4-3* | Est-1 | c.14_50del | p.(Gly5Alafs*57) |
| *rpb1-like4-4* | Est-1 | c.15_363del | p.(Ser6Tyrfs*19) |
| Name | Background | Impact on *RPB1* and *RPB1-like-4* genes | Impact on RPB1 and RPB1-like-4 proteins |
| *rpb1-5 rpb1-like4-5* | Uod-1 | *RPB1*:c.47_418del<br>*RPB1-like-4*:c.14del,<br>c.363_364insT | RPB1:p.(Gly16Alafs*25)<br>RPB1-like-4:p.(Gly5Valfs*12) |

The Uod-1 line carrying mutations in both *RPB1* and *RPB1-like-4* was susceptible to clubroot infection but did not exhibit any enhanced symptoms compared with Col-0 or the single mutants for *RPB1*. The development of clubroot disease in *rpb1* and *rpb1-like-4* mutants was further characterised by microscopic observation of longitudinal sections of the hypocotyl and upper root 25 dpi when the pathogen life-cycle will have progressed to the formation of resting spores. The expansion of galls was even more prominent in the *rpb1* mutants at this time with the extensive swelling of hypocotyls and roots and reduction in fine roots (Supp Fig. 7B), whereas in the resistant wildtype Est-1 and Uod-1 the only symptoms of infection that could be observed were some small galls formed on secondary roots and some darkening of patches of the roots that could be due to cell wall lignification or other defence responses. Longitudinal sections of these galls revealed in the *rpb1* mutants, as in Col-0, an abundance of enlarged host cells filled with secondary plasmodia and resting spores (Figure 4, Supp. Fig. 8). The vasculature of these plants was disorganised with reduced xylem whereas in wildtype Est-1 and Uod-1 the xylem was contiguous and as abundant as uninfected controls, the only signs of infection were isolated plasmodia restricted to the epidermis often surrounded by highly lignified cells (Figure 4, Supp Fig. 9).

**Figure 4.**
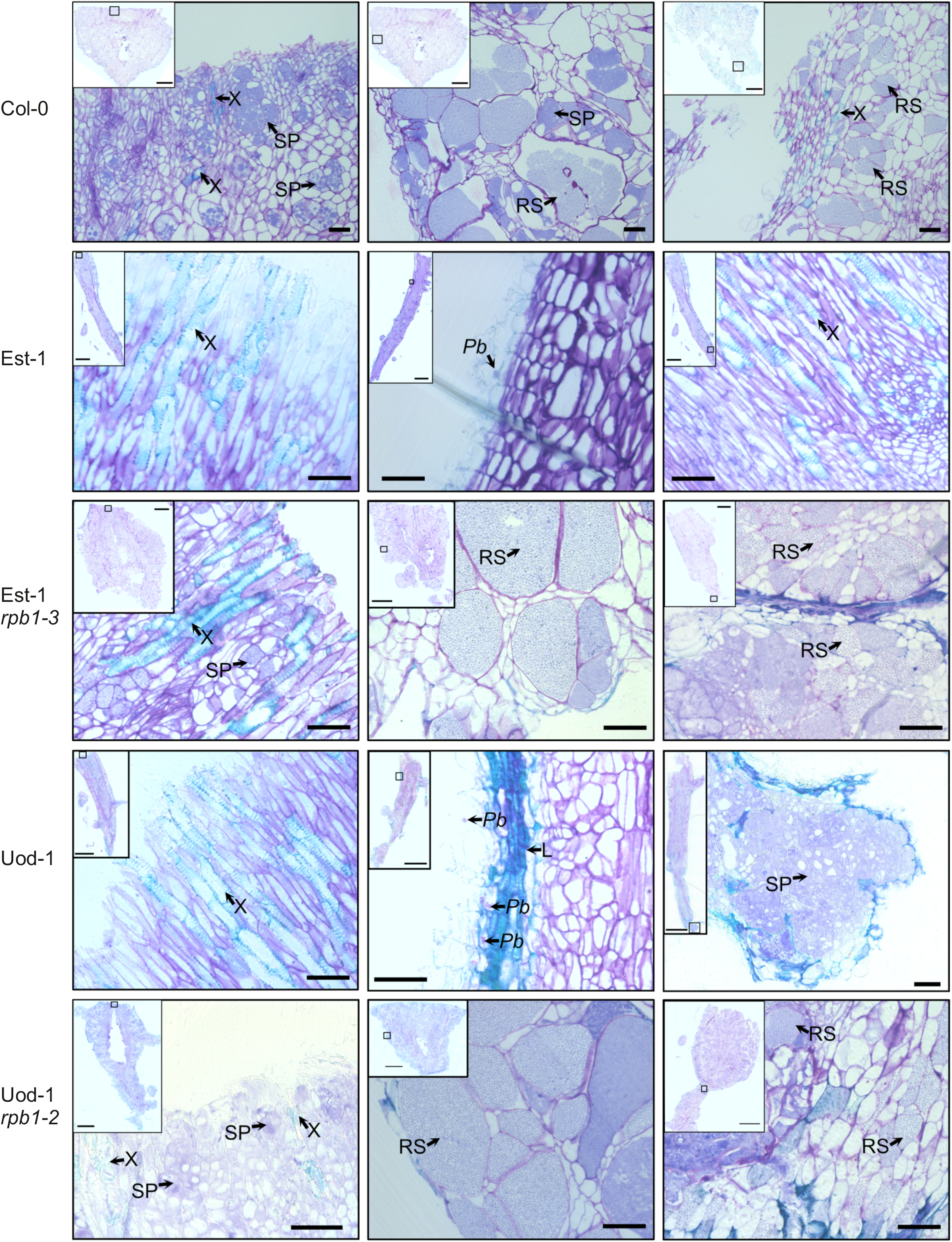
Clubroot disease development in *rpb1* mutants is comparable to susceptible Col-0. Histological characterisation of the hypocotyls of wildtype accessions and *rpb1* mutants inoculated with *P. brassicae* 25 dpi. Micrographs in the left column focus on symptoms in the uppermost central part of the hypocotyl, the central column focusses on the epidermal cells. In the right column the region of the interphase between the hypocotyl and the root is presented. X: Xylem cells, L: Lignified tissue, RS: Resting spores, SP: Secondary plasmodia, Pb: *P. brassicae* cells. The scale bar in the inset overview picture corresponds to 1000 μm, the scale bar in the detailed pictures corresponds to 50 μm.

The activation of defence signalling in response to *P. brassicae* was assessed at the much earlier time-point 7 dpi, which represents the early stages of secondary infection when secondary zoospores attempt to colonise the central root cortex. Expression of the defence marker gene *PR5* (Lemarié et al., 2015a), the enzyme critical for phytoalexin biosynthesis *CYP71A13* (Nafisi et al., 2007), and the defence signalling component *AED1* (Breitenbach et al., 2014), are all significantly upregulated in response to infection in Est-1 and Uod-1 at this time, while *PR5* is also upregulated in the susceptible Col-0 although by a smaller amount (Figure 5B, C, D). The loss of *RPB1* significantly reduced the activation of these genes with *PR5* being upregulated to a reduced degree in the Uod-1 *rpb1-2* mutant while in other cases expression was the same as in mock controls. Interestingly, though when the expression of *RPB1* was examined (using primers targeting transcript regions outside of the deletions) there was significant activation of the *RPB1* promoter in the Est-1 *rpb1-3* mutant in response to infection and *RPB1* transcript could be detected in the Uod-1 *rpb1-2* only with *P. brassicae* inoculation, which, in this experiment, was the same case for wildtype Uod-1 (Figure 5A).

**Figure 5.**
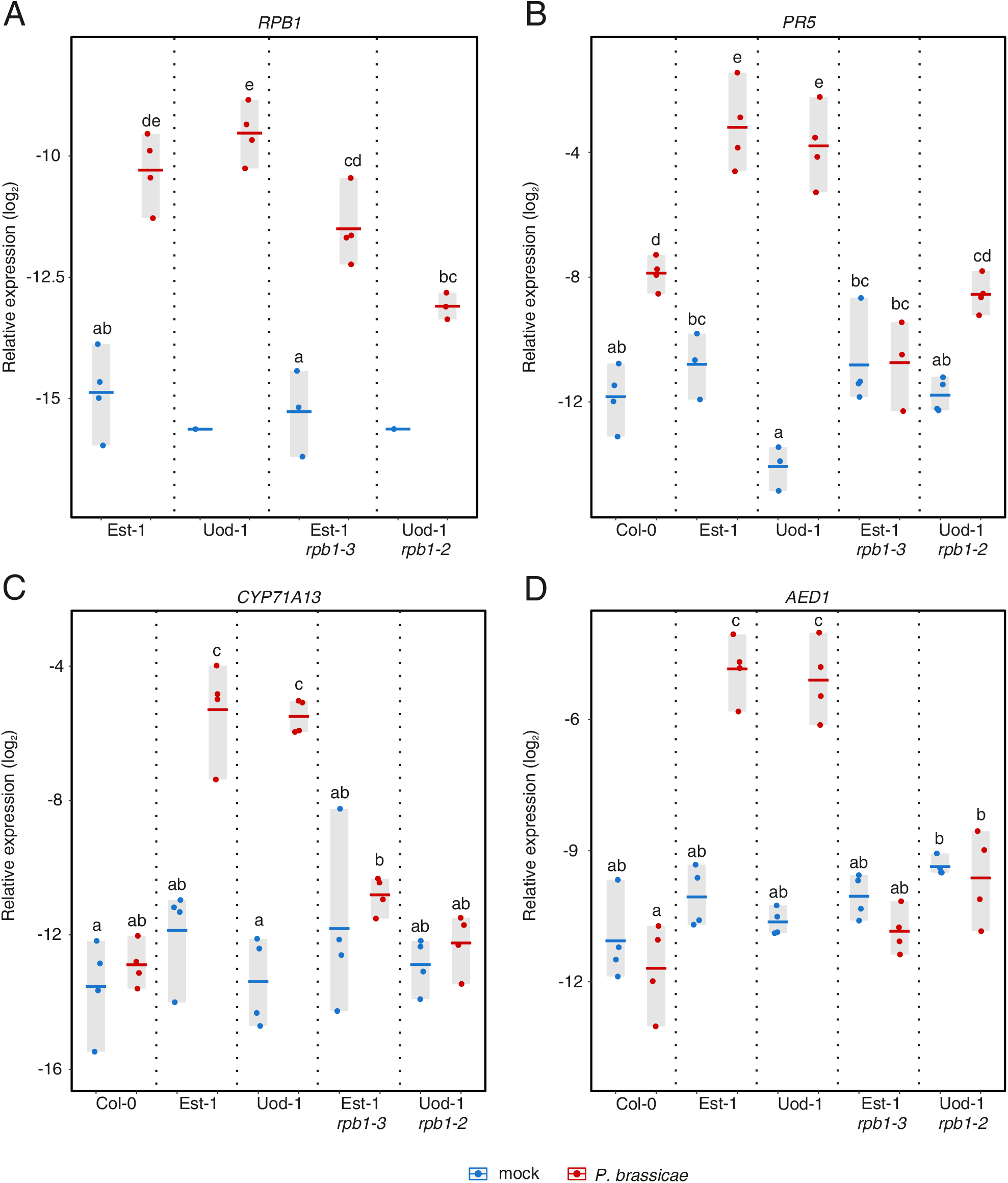
Defence responses to *P. brassicae* in Est-1 are dependent on *RPB1*. Expression of A) *RPB1* transcript, B) *PR5*, C) *CYP71A13* and D) *AED1* in *P. brassicae* inoculated *rpb1* mutants 7 dpi. Expression is relative to *AT1G76030* and *AT3G48140*. Different letters indicate statistically significant differences (BH adjusted p < 0.05). Each point represents one biological replicate (n = 4) of 10 plants, horizontal bars indicate the mean. *RPB1* transcript was below the limit of detection in 3 of the 4 biological replicates in the mock inoculated Uod-1 plants.

### Ectopic expression of *RPB1* in a susceptible background is not sufficient for resistance to clubroot

*RPB1* encodes a protein with no homology to previously characterised proteins, it is predicted to contain membrane spanning regions (Supp. Fig. 10) but no known protein domains are present. Orthologues of *RPB1* can be found in the Brassicaceae and there is a homologue of *RPB1* in the Col-0 genome – *AT1G56270*, however there is no information available regarding the role or function of these related genes (Supp. Fig. 11). To begin the characterisation of *RPB1* the gene was cloned from Est-1 and placed under the control of the *CaMV35S* promoter for over-expression studies. Transient expression of *35S::RPB1* via agrobacterial transfection of *Nicotiana tabacum* leaves induced cell death after 72 hours however this was not observed with N-terminal fusion of GFP to the RPB1 protein (Figure 6A). No transformants could be obtained in Arabidopsis accession Col-0 with the *35S::RPB1* construct despite repeated efforts. Transformation of Col-0 with *35S::GFP-RPB1* produced plants with a wide range of growth restriction phenotypes with some transformants appearing close to wild-type and others being too small to produce seed. T3 progeny of three independent transformants were assessed for their response to *P. brassicae*. Only plants that were considered viable in terms of size at the time of inoculation were included, removing some of the most extreme examples of growth restriction. At 19 dpi the expression of *GFP-RPB1* was determined in the leaves of transformants and the symptoms in the galls were assessed. *GFP-RPB1* expression was found to have a significant negative correlation with clubroot symptoms for both hypocotyl expansion (Spearman Rho = −0.83) and relative *P. brassicae* DNA levels (Spearman Rho = −0.61) (Figure 6B, C). Plant size as determined by rosette diameter was also significantly negatively correlated with *GFP-RPB1* expression in the leaves (Spearman Rho = −0.76) (Figure 6D). Expression of the defence / stress marker gene *PR-1* was measured in the leaves and found to be positively correlated with *GFP-RPB1* expression (Spearman Rho = 0.66) (Figure 6E). With constitutive expression of *RPB1* in Col-0 having the potential to cause such dramatic effects on growth the native promoter of *RPB1* from a resistant accession was assessed. *RPB1* and the 1024 bp upstream promoter region was cloned from Est-1; five transgenic Col-0 lines were obtained which did not exhibit any restriction in growth and were morphologically similar to wildtype Col-0, these were assessed for their response to *P. brassicae* infection 19 dpi. Two lines (6 and 9) were indistinguishable from wildtype Col-0 in terms of gall width and pathogen DNA levels (Figure 7B, C) despite the fact that *RPB1* expression was induced in response to infection at 7 dpi indicating that the *RPB1* promoter could be activated in a Col-0 background (Figure 7E). None of the lines were fully resistant to clubroot or comparable to Est-1 however lines 1, 4, and 7 had significantly lower accumulation of *P. brassicae* DNA and line 1 had significantly smaller galls (Figure 7A, B, C, D). While expression of *RPB1* in lines 4 and 7 was also induced in response to *P. brassicae* at 7 dpi there was no up-regulation of *RPB1* in line 1, rather in this line the expression of *RPB1* was constitutively high matching the level of expression in *P. brassicae* inoculated Est-1 in both the mock-treated and infected samples (Figure 7E). Only *pRPB1::RPB1* line 1 responded to infection with the activation of defence genes *CYP71A13* and *AED1* (Figure 7F, G). Thus, it appears that clubroot susceptible accession Col-0 has the capacity to trigger partially effective responses to *P. brassicae* infection dependent on the level, or possibly timing, of *RPB1* expression.

**Figure 6.**
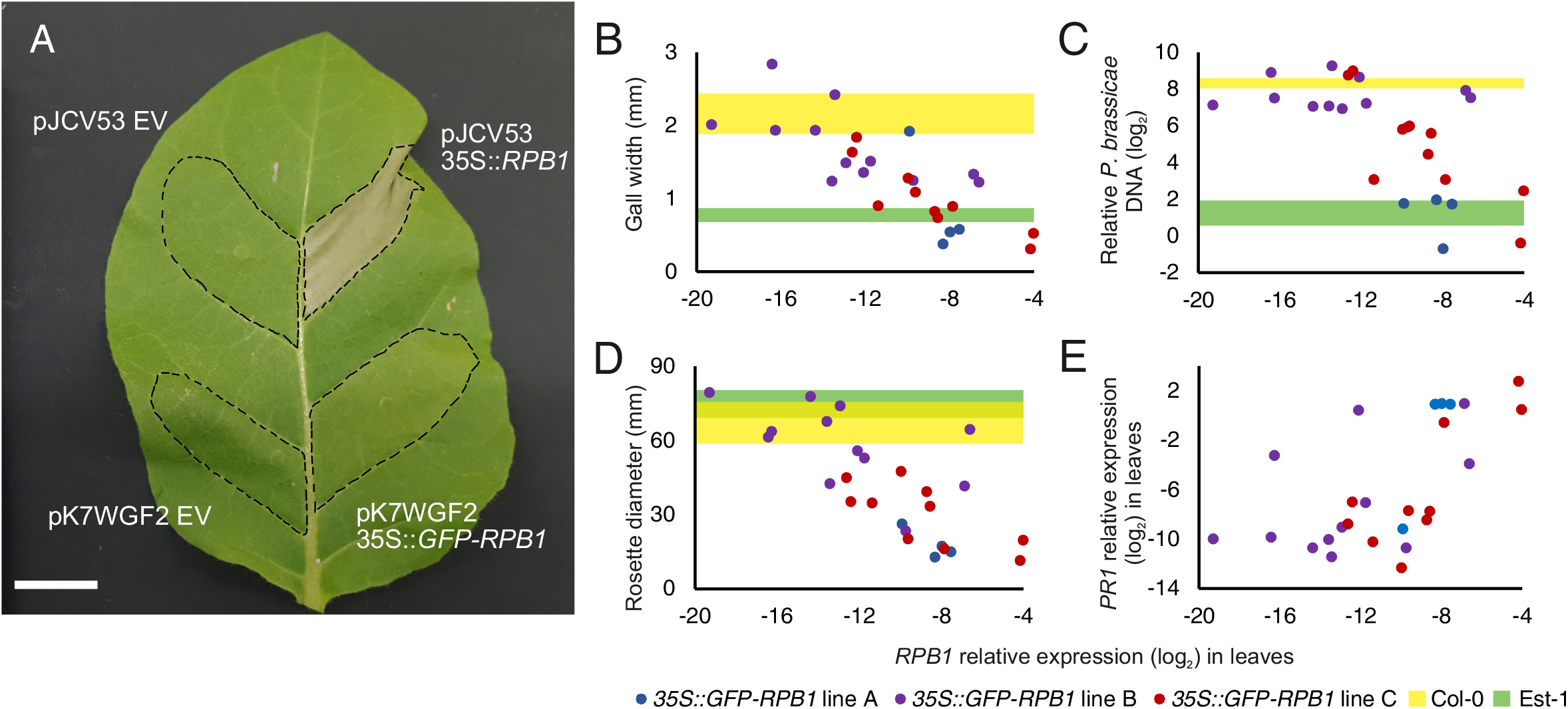
Ectopic over-expression of *RPB1* induces stress responses. A) Transient expression of *35S::RPB1* and *35S::GFP-RPB1* in *N. tabacum* leaf three days after agroinfiltration. Scale bar = 2 cm, EV empty vector. B-E) Expression of *RPB1* (relative to *AT1G76030, AT3G18780* and *AT3G48140)* in the leaves of Col-0 *35S::GFP-RPB1* transformants 19 dpi plotted against: B) gall width; C) relative pathogen DNA titre (*Pb18S/AtSK11*); D) rosette diameter; E) relative *PR-1* expression in the leaves. Individual points represent pools of 3-6 T3 plants with a common T2 parent, coloured bands for Col-0 (yellow) and Est-1 (green) represent the 95% confidence interval for 15 wildtype plants (n = 5 for DNA quantification).

**Figure 7.**
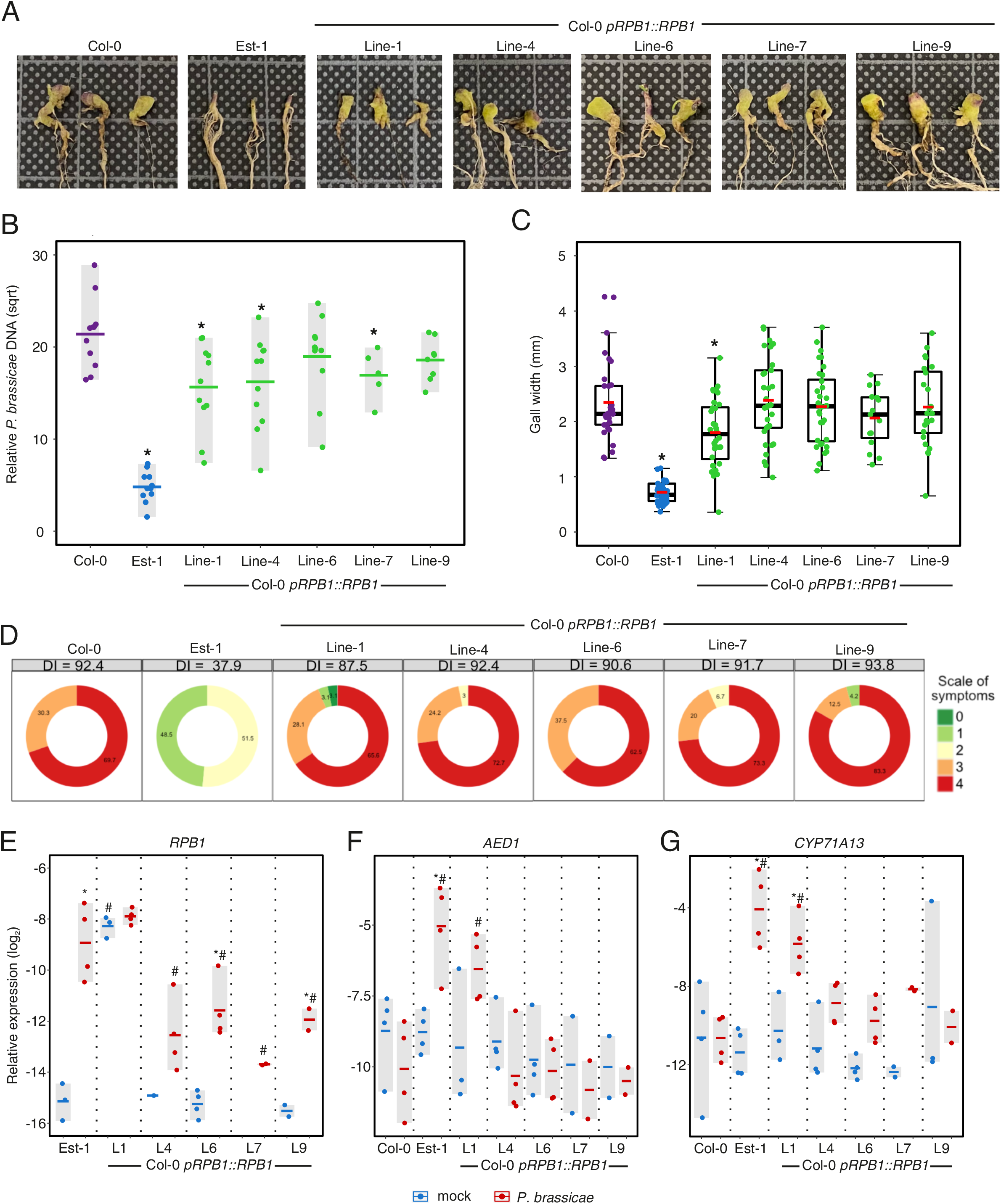
Expression of *RPB1* in Col-0 does not confer clubroot resistance. A) Gall symptoms in Col-0 *pRPB1::RPB1* transgenic lines 19 dpi. Squares on the background grid have 10 mm length. B) Relative pathogen DNA titre (*Pb18S/AtSK11*) 19 dpi, points indicate biological replicates of 3 plants, horizontal lines indicate the means, asterisks indicate statistically significant differences from wildtype Col-0 (Tukey, p < 0.05), 5 ≤ n ≤ 11. C) Gall width measurements for individual plants pooled for B, red lines indicate the mean, black boxes the lower quartile, median and upper quartile, whiskers the 1.5 IQR, 15 ≤ n ≤ 33, asterisks indicate statistically significant differences from wildtype Col-0 (Games-Howell, p < 0.05). D) DI score for the symptoms of galls in B & C, the percentage of plants assigned to each symptom class is shown on each segment. E-G) Expression of *RPB1, AED1* and *CYP71A13* in wildtype Col-0, Est-1 and Col-0 *pRPB1::RPB1* transgenic lines 7 dpi (relative to *AT1G76030* and *AT3G48140*). Each point represents one biological replicate (2 ≤ n ≤ 4) consisting of the roots and hypocotyls of 6 plants, horizontal bars indicate the mean. *RPB1* transcript was below the limit of detection for some replicates in mock inoculated lines 4, 7 and 9. Asterisks indicate statistically significant differences from mock treatment while hash marks indicate statistically significant differences from wildtype Col-0 (F & G) or Est-1 (E) (p < 0.05).

## DISCUSSION

### Natural variation in Arabidopsis responses to *P. brassicae*

The *P. brassicae* isolate used in the study represents a pathotype that is particularly devastating and is capable of infecting oilseed rape cultivars that carry the ‘Mendel’ resistance locus (Řičařová et al., 2016; Ramzi et al., 2018; Zamani-Noor et al., 2022). Its aggressive nature was born out in the assay of 142 Arabidopsis accessions, moderate to strong symptoms of clubroot disease were established in all but 11 of the accessions. Within the susceptible accessions there was a range in the severity of infection whether quantified by relative pathogen DNA levels or by inspection of gall development. However, GWAS did not identify any SNPs that were significantly correlated with the degree of infection when analysis was restricted to the clubroot susceptible lines. The progress of clubroot disease within susceptible accessions is likely governed by complex multi—gene interactions and it may be that a much larger panel of accessions would be needed in order to provide enough statistical power to identify the key components. Another consideration in refining this study would be to consider the impact of differences in host flowering time. The relatively slow progress of clubroot disease in Arabidopsis means that, by the time symptoms have developed to the point where they can be reliably scored, some early flowering accessions may have already bolted; the impact of flowering on distribution of host resources will have knock on effects on the availability of nutrients being delivered to galls and so significant associations may be uncovered in the analysis of cohorts of accessions with matched flowering times, which has been the case for GWAS analysis of the interaction between *Botrytis cinerea* and Arabidopsis (Corwin et al., 2016). For at least one accession (Var2-6) the effect of *P. brassicae* infection was more apparent in the aboveground tissue than in the development of galls in the root system, hyper-susceptible accessions may prove so sensitive to the disruption of vascular tissue wrought by *P. brassicae* that symptoms of drought curtail gall development. An integrated analysis of above and below-ground symptoms may have the added benefit of identifying accessions with greater tolerance of infection and mitigation of water deficit stresses associated with pathogen manipulation of xylem and phloem development.

GWAS analysis of the full complement of accessions identified three loci that correlated with resistance. The variation in sequences upstream of *WIP2* are a potential focus for further investigation, since the deletions and polymorphisms in the promoter region may affect the functioning of *WIP2*, which was found to have higher transcript levels in susceptible Col-0. While *WIP2* expression was not up-regulated in response to infection at the 7 dpi time-point evaluated (Supp. Fig. 3B), in RNA-Seq profiling of Col-0 roots responding to *P. brassicae* infection 16 dpi *WIP2* is significantly down-regulated while three other members of the five gene family (*WIP3, WIP4, WIP5*) are up-regulated (Rolfe et al., 2016). Two orthologues of these *WIP* genes from soybean were found to reduce susceptibility to *Pseudomonas syringae* pv tomato DC3000 when heterologously expressed in Arabidopsis. The potential of these peptides to restrict growth of biotrophic pathogens and the specific suppression of *WIP2* at later stages of *P. brassicae* infection may warrant further investigation of their role in responses to clubroot disease.

### *RPB1* is essential for clubroot resistance in Est-1 and Uod-1

Of the loci implicated in resistance to *P. brassicae* that were tested through the generation of knock-out mutations only the gene *RPB1*, the locus of which was previously identified by Fuchs and Sacristan (1996) and sequenced by Rehn and Siemens (unpublished), was confirmed to impact clubroot resistance. The deletions of *RPB1* in accessions Est-1 and Uod-1 accumulated pathogen DNA titres that were comparable to the highly clubroot susceptible accession Col-0 19 dpi (Figure 3B). The average DI score for *rpb1* mutants in Est-1 and Uod-1 were 88.6 and 93.4 (Figure 3D); when ranked by DI score the wildtype accessions were positioned 142^nd^ and 139^th^ out of 142 accessions tested while the *rpb1* mutant DI scores would positioned 32^nd^ and 7^th^. The ability of *P. brassicae* to colonise the cortex of *rpb1* mutants and develop mature resting spores 25 dpi was indistinguishable from Col-0. Despite the sequence similarity of RPB1-like-4 (56.7% aa similarity) and its transcriptional activation in response to *P. brassicae* no impact on clubroot disease development was determined in the *rpb1-like-4* mutants, furthermore no exacerbation of symptoms was observed in the *rpb1 rpb1-like-4* double mutant indicating that RPB1 plays a specific and essential role in resistance to *P. brassicae*.

### Potential roles for RPB1 in mediating clubroot resistance

RPB1 represents a novel plant defence response component. While it has several orthologues among the Brassicaceae it has no known molecular function and no similarity to previously characterised proteins. Lacking any of the domains typically associated with recognition of non-self it is unlikely that RPB1 could act alone to mediate the recognition of a *P. brassicae* avirulence factor. *RPB1* expression is activated even in the absence of a functional RPB1 protein, though not to the same degree as in wildtype Est-1 (Figure 5A). Ectopic expression of RPB1 alone in *N. tabacum* can induce a hypersensitive response (HR) while over-expression in Col-0 is either unviable or highly deleterious even for a potentially mis-folded GFP-RPB1 fusion protein (Figure 6A, D). The transmembrane domains predicted for RPB1, its size and capacity to induce HR do bear some resemblance to the group 2 executor genes identified in rice and pepper (Ji et al., 2022). Executor proteins, which are activated by the TAL effectors of *Xanthomonas* spp., do not have conserved sequences but share some structural features, the group 2 E proteins are characterised by multiple membrane spanning domains with (Leu / Ile) – X (4-9) – (Leu / Ile) motifs, RPB1 is predicted to contain four such domains with Leu / Ile motifs as do XA10 from rice and Bs4C-R from *Capsicum pubescens*. Another feature is the presence of Glu and Asp residues in the C-terminal region, while no Asp residues are present in this part of the RPB1 sequence there is one Glu residue, the same as Bs4C-R. While these similarities are somewhat tenuous it would be interesting to confirm whether the presence of the C-terminal acidic residue is essential for HR elicitation in non-host species as is the case for executor proteins. Similarly, if an appropriate RPB1 fusion protein, capable of fluorescence, can be obtained localisation to the ER membrane would be more evidence for potential similarity to this class of resistance proteins. The other potential role for RPB1 would be in the elaboration of an immune response signal through the interaction with other factors; while *RPB1* expression is activated in response to *P. brassicae* when the gene and promoter from Est-1 are transferred to Col-0 there is no equivalent clubroot resistance conferred indicating that other factors are not functioning. The activation of downstream defence responses such as the biosynthesis of phytoalexins via CYP71A13 (Nafisi et al., 2007) or the elaboration of systemic acquired resistance via AED1 (Breitenbach et al., 2014) were only potentiated when the transgene copy number or position effects specific to Line-1 acted to drive *RPB1* expression in mock treated transgenic plants to equivalent levels as the *P. brassicae* inoculated wildtype Est-1 (Figure 7E, F, G). The magnitude, timing and perhaps localisation of *RPB1* expression may be critical to its effective function in mediating resistance but even *pRPB1::RPB1* Line-1 with its constitutive activation was significantly more susceptible to clubroot disease than wildtype Est-1 underlining the potential requirement for additional factors absent in Col-0.

Inspection of sections from Est-1 and Uod-1 *P. brassicae* infected root systems revealed that colonisation by the pathogen was restricted to the epidermis with the neighbouring cells surrounding an infection site typically forming thickened, highly-lignified cell walls (Supp. Fig. 9). While the interactions between host and pathogen may be restricted to a small number of cells in the epidermis in resistant accessions the measurements for changes in gene expression come from whole root tissue samples, it would be very interesting to determine any *RPB1* dependent defence gene expression responses at these infection sites, compared to any global responses in the roots. Expression of *RPB1* appears to be under tight control, in several mock inoculated plants the transcript could not be detected by qPCR. While the aa sequence of RPB1 protein is conserved in the clubroot susceptible accessions that carry it (C24, Cvi-0, Kyoto, An-1) the upstream promoter sequence appears to be conserved only in the resistant accessions. Sequencing of this region in additional resistant and susceptible accessions may shed light on any conserved cis-elements that are involved in the regulation of *RPB1* expression.

### Expression patterns of *RPB1* related genes

The only RPB1 related gene to be examined so far is *RPB1-like-4*, which is not required for resistance to *P. brassicae*. While Col-0 lacks an *RPB1* gene one homologue, *AT1G56270*, was identified. Inspection of *AT1G56270* expression patterns in publicly available microarray datasets (Zimmermann et al., 2004) did not reveal any significant responses to pathogen treatment or with biotic interactions; rather the highest expression levels and greatest differential expression were in pollen compared with other tissue types, indicating that this *RPB1* homologue may be involved in tissue specific or developmental functions (Pina et al., 2005, E-MEXP-285). Six orthologues of *RPB1* can be identified in the genome of *B. napus* cultivar Darmor-bzh (Supp. Fig. 11). It is interesting that *RPB1* orthologues have so far not been identified in any of the screens for clubroot resistance in *B. napus* or its parental species *B. rapa* and *B. oleracea*. Recently, RNA-Seq profiling of responses to *P. brassicae* in clubroot resistant *B. napus* oilseed rape and rutabaga cultivars have been published (Galindo-González et al., 2020; Zhou et al., 2020). Mapping of these reads (PRJNA597078, PRJNA641167) to the Darmor-bzh genome revealed no differential expression in response to infection, or between resistant and susceptible genotypes for any of the putative *RPB1* orthologues, in fact no transcripts were detected for three genes while relatively low, stable expression levels were observed for the other three. Since the majority of publicly available gene expression data for Arabidopsis is derived from the RPB1-lacking Col-0 background it will be interesting to identify the conditions under which the orthologues most closely resembling *RPB1* from Brassica species are transcriptionally activated.

### Possible role for RAC1 in clubroot interactions

The pseudogenisation of *RPB1* in Col-0 combined with the potential of RPB1 to induce HR in *N. tabacum* and GFP-RPB1 to restrict growth in Col-0 indicates that RPB1 protein (or proteins resembling RPB1) imposes fitness penalties on genotypes where its expression is not tightly controlled. The association of polymorphisms in the sequence of resistance gene *RAC1* with accessions that were resistant to clubroot disease initially lead us to hypothesise that RAC1 could be involved in recognition of some *P. brassicae* avirulence factor however *rac1* mutants remained completely resistant to clubroot disease. *RAC1* is a hotspot for meiotic crossover leading to sequence diversification (Choi et al., 2016). It may be that the association of RAC1 polymorphisms with clubroot resistance are a statistical artefact due to the small sample size, it is interesting to speculate that while some alleles of *RAC1* are involved in the recognition of non-self (Borhan et al., 2004), in other accessions RAC1 could possibly be involved in the monitoring or regulation of RPB1 protein expression to mitigate any fitness costs implicated in the clubroot resistant accessions where it has not be pseudogenised and changes to its promoter sequence, or other epigenetic factors, have not limited its expression to an even greater degree.

### Future directions for further elucidation of the mode of action of RPB1 in model and crop plants

The search for putative protein-protein interactions between RPB1 and other host factors will be an important step for the further characterisation of this critical component of the clubroot resistance response in Arabidopsis. Identification of regulators involved in its transcriptional activation in resistant accessions may also prove key. The failure of *RPB1* expression to provide clubroot resistance in Col-0 points to the requirement for either additional components or appropriate expression dynamics. Determining these factors may provide a strategy for utilising *RPB1* for clubroot resistance in crop brassicas either through augmentation of endogenous host resistance machinery, via co-transformation with accessory proteins from Arabidopsis or through the identification of appropriately *P. brassicae*-responsive promoter elements for crop cultivars. Establishment of the subcellular localisation of RPB1 and its potential for membrane binding will likely also shed light on potential function. RPB1’s capacity to elicit PCD in *N. tabacum* offers one avenue to test the importance of various residues in the protein sequence such as those shared with rice executor proteins or serine residues predicted to be potential sites of phosphorylation and whether they are required for immune functioning. RPB1 is required for the full activation of certain defence genes that are *P. brassicae* induced even in clubroot susceptible backgrounds (*PR5*) or solely in resistant accessions (*AED1, CYP71A13*). Transcriptional profiling of RPB1 dependent responses to *P. brassicae* infection may indicate which subset of clubroot responsive genes are critical for the mounting of a successful defence. The clubroot resistance accessions described here have proved resistant to the other *P. brassicae* pathotypes collected from Poland tested so far; in the original identification of *RPB1*, Tsu-0 was reported to be susceptible to an isolate collected from a *Brassica napus* cultivar used in the European clubroot differential set ECD07 (Buczacki et al., 1975; Fuchs and Sacrisatan, 1996). In order to dissect the function of RPB1 in mediating resistance responses it will be useful to characterise responses in a compatible interaction.

## MATERIALS AND METHODS

### Plant material and growth conditions

Arabidopsis accessions were obtained from the Nottingham Arabidopsis Stock Centre (NASC) the seed stock identifiers are listed in Supp. Table 1. Plants were grown in controlled conditions with a short-day photoperiod (9 h light / 15 h dark; 22 °C / 20 °C). FluorA L 36W/77 lamps with an irradiance of 120 μmol m^-2^ s^-1^ were used and a relative humidity of 65% was maintained in the chamber. For the screen of accessions plants were sown directly on soil following stratification for 4 days at 4 °C degrees in distilled water. For the evaluation of transgenic lines, seeds were surface sterilised with bleach and germinated on Murashige and Skoog (MS) medium agar plates for 10 days prior to transfer to soil. Plants were grown on a 5:1 mixture of Klasmann No. 11 soil (pH 6.3) and perlite, except when roots were harvested for RNA extraction when a 1:1 sand soil mix was used.

### Pathogen propagation and inoculation

The *P. brassicae* isolate used was classified as pathotype P1 according to the Somé differential set (Somé et al., 1996), additionally it was observed to infect the clubroot resistant *B. napus* cultivar ‘Mendel’, and was designated as P1+ (Zamani-Noor et al., 2021). The Chinese cabbage *B. rapa* var. pekinensis cultivar ‘Granaat’ was used for pathogen propagation; pathogen spores were prepared according to the method of Fuchs and Sacristan (1996), and spore concentration determined with a haemocytometer. A spore concentration of 1 x 10^6^ spores ml^-1^ was used, each Arabidopsis plant was inoculated by pipette with 2 ml of spore suspension at 17 days after sowing.

### Clubroot phenotype scoring

The hypocotyl and upper 1 cm of root were harvested from plants 19 dpi and photographed for symptom scoring and gall measurement. DNA extraction was performed from the combined tissue of three plants. Tissue was macerated in DNA extraction buffer (100 mM TRIS-HCl pH 8.0, 50 mM EDTA pH 8.0, 500 mM NaCl, 1.3% SDS) in a TissueLyser II (Qiagen, Germany) with two metal beads for 2 minutes at 30 Hz. DNA was purified with KAc and precipitated with isopropanol and NaAc, followed by RNAse treatment (Qiagen, Germany) and subsequent re-precipitation. For quantitative PCR 90 ng of template DNA was used in 10 μl qPCR reactions with 0.25 μM primers and Luna qPCR Master Mix (NEB, USA). qPCR was performed in a LightCycler 480 (Roche, Germany), system software was used to calculate Cp values according to the 2nd derivative max method for host target *AtSK11 (At5g26751*) and pathogen target *Pb18S* (ENSRNAT00050137123), primer sequences are listed in Supp. Table 3. Two technical replicates were performed for each gene for each sample and were averaged; the pathogen DNA titre relative to the host was calculated from difference between Cp*Pb18S* and Cp*AtSK11*. Clubroot disease index scoring was performed according to the method of Ludwig-Müller et al., (2017), using the following scale: 0 - no symptoms observed; 1 - presence of small clubs only in lateral roots, root architecture is preserved; 2 - small clubs present in the main and lateral roots with thickening of the main root but no visible symptoms in the hypocotyls; 3 - medium to large clubs and galls are observed in the main and lateral roots, most of the fine roots are absent and the hypocotyls exhibits some swelling; 4 - severe swollen galls in the roots and hypocotyls reaching the rosette. The DI score was calculated as a percentage from the formula (1n_1_+ 2n_2_+ 3n_3_+ 4n_4_)/4N, where N is the total number of plants evaluated and n_1_ to n_4_ denote the number of plants assigned to each clubroot gall class. Gall width measurements from photographs were made using ImageJ software, taking a subjective assessment of the maximum gall diameter perpendicular to the growth of the hypocotyl or root (Schneider et al., 2012).

### Genome-wide association analysis

Where necessary, normalisation of the accession phenotype data was performed using the bestNormalize package in the R environment (Peterson, 2021). GWAS was performed using two online toolsets: GWA-Portal (https://gwas.gmi.oeaw.ac.at/) and easyGWAS (https://easygwas.ethz.ch/) (Seren et al., 2013; Grimm et al., 2017) analysing subsets of available genotype data with the AMM and EMMAX algorithms respectively. Genotype data for accessions in these platforms come from the 1001 Genome sequencing project and the RegMap panel of accessions genotyped with the Affymetrix 250k chip (Horton et al., 2012; Alonso-Blanco et al., 2016).

### Generation of CRISPR/Cas9 knock-out mutations

The strategy used for introducing mutations to target genes was previously described by Bieluszewski et al., (2022); two target sequences spaced 200 – 400 bp apart were selected to generate polymorphisms that could potentially be identified by PCR. Guide RNAs (gRNAs) were designed using the CRISPOR tool, to evaluate the potential likelihood of generating double strand breaks while predicting possible off-target binding sites in the genome (Concordet and Haeussler, 2018). The selected gRNA spacers (matching genomic targets) were chemically synthesised and cloned by PCR into the pJET1.2 based gRNA vectors using Clontech CloneAmp HiFi PCR Premix (Takara, Japan). The transformation vector pICU2:Cas9-dsRED containing the *Cas9* gene from *Streptococcus pyogenes* expressed under the promoter of the Arabidopsis *Incurvata 2* gene (*ICU2, At5g67100*) was combined with the gRNA combinations by Gibson assembly (NEB, USA). For *RPB1, RPB1-like-4*, and *RAC1* targeting, two gRNAs were incorporated; for the combined targeting of *RPB1* and *RPB1-like-4* four gRNAs were incorporated into the vector. The construct was transformed into *A. tumefaciens* EHA105 which was used to transform Arabidopsis accessions by the floral dip method (Clough and Bent, 1998). Transformed seeds were identified by dsRED fluorescence under an AxioZoom V16 monoscope (Zeiss, Germany). In the subsequent generation non-fluorescent T1 seeds were selected for the screening of stable mutations in the absence of the Cas9 gene. Pools of plants were genotyped using the Phire Tissue Direct PCR Master Mix (Thermo, USA) and primers flanking the targeted regions, from these pools individual mutants were selected and homozygous mutants identified in the subsequent generation were confirmed by sequencing. All primers used for cloning and genotyping are detailed in Supp. Table 3.

### *RPB1* cloning

*RPB1* was cloned from Est-1 as the (1) coding sequence (CDS) including the stop codon and (2) the CDS and native promoter region 1024 bp upstream using Phusion High Fidelity DNA polymerase (NEB, USA) and TA-ligated into the entry vector pCR8/GW/TOPO (Thermo, USA). Construct sequences were confirmed by Sanger sequencing and were cloned into destination vectors (1) pJCV53 and pK7WGF2 and (2) pKGWFS7 via LR reactions to make (a) pJCV53-*35S::RPB1*, (b) pK7WNGF2-*35S::GFP-RPB1* and (c) pKGWFS7-*pRPB1::RPB1*. These destination constructs were transformed into *A. tumefaciens* EHA105 for transient expression in *N. tabacum* cv Xanthi and floral dip transformation of Arabidopsis accession Col-0 (Clough and Bent, 1998; Norkunas et al., 2018). Transgenic Col-0 plants were identified on MS agar plates supplemented with 50 μg m^-1^ kanamycin. In experiments performed on segregating T2 and T3 lines individual plants were genotyped 14 days after germination using the Phire Tissue Direct PCR Master Mix (Thermo, USA). The primers used for cloning and genotyping are listed in Supp. Table 3.

### Light microscopy

Tissue for microscopy was collected in a fixative solution containing 0.5% glutaraldehyde and 2% paraformaldehyde in PBS buffer pH 7.4 and incubated at 4 °C for 3-6 days. Samples were then transferred to 10% ethanol for at least 30 min, and then subsequently the ethanol concentration was increased successively to 30%, 50%, 70% and absolute ethanol for intervals of 30 min and then preserved in absolute ethanol for at least 2 days. Samples were then preinfiltrated with a solution of 1:1 ethanol/Technovit 7100 (v/v) and ultimately infiltrated with 100% Technovit 7100 (Kulzer, Germany) (Stefanowicz et al., 2021). The sectioning of the embedded samples was performed on a RM2135 microtome (Leica, Germany) at a thickness of 5 μm and transferred to glass slides with a drop of distilled water and dried on a heating plate at 60 °C. Sections were stained using 0.05% Toluidine Blue (w/v) solution. Photographs were taken with an Axio Image M2 microscope coupled to an AxioCamICc5 camera (Zeiss, Germany).

### Gene expression analysis

For RNA extraction from root samples, all root material was collected along with the hypocotyl and quickly washed before freezing in liquid N_2_. RNA was extracted from pulverised tissue using the InviTrap Spin Plant RNA Mini Kit (Invitek, Germany) using the DCT buffer and β-mercaptoethanol. RNA was quantified using by Nanodrop 2000 (Thermo, USA), 1 μg RNA was treated with TURBO DNAse (Thermo, USA) and first strand cDNA was synthesised with M-MLV Reverse Transcriptase, RNase H Minus (Promega, USA). qPCR was used to quantify relative transcript levels using Luna qPCR Master Mix (NEB, USA) in the LightCycler 480 instrument (Roche, Germany). The system software calculated Cp values according to the 2nd derivative max method for genes of interest relative to expression levels of the normalisation genes *AT1G76030 (VAB1), AT3G48140* and *AT3G18780* (*ACT2*). For each sample two technical replicates were run and their values averaged, expression relative to the geometric mean of two or three normalisation genes was then calculated.

### Bioinformatic and statistical analysis

Analysis of gall width measurements and qPCR data for relative pathogen DNA titre in knockout mutants and transgenic lines was performed using linear models in the R environment using the package rstatix (R Core Team, 2022). For the qPCR data of knock-out and transgenic lines to satisfy the requirements of linear models for normality and homoscedasticity it was necessary to transform the data by taking the square root of the un-logged Cp values. For the analysis of gene expression data mixed-effect linear models were applied to the Cp values taking genotype and treatment as fixed effects and experiment as random effect using the lme4 package in R (Bates et al., 2015). When multiple testing correction was appropriate to control the false discovery rate the Benjamini and Hochberg (1995) correction was used, while the Bonferroni correction was applied in the GWAS analysis (Bland and Altman, 1995). Prediction of membrane spanning regions in the RPB1 amino acid sequence were made using PHOBIUS (Käll et al., 2004). Orthologues of RPB1 were identified using Orthofinder (Emms and Kelly, 2019) to analyse the proteomes of Arabidopsis TAIR10 Col-0 (+ RPB1 and RPB1-like sequences), various Brassica species from the BRAD database (Chen et al., 2022), *B. napus* Darmor-bzh (Rousseau-Gueutin et al., 2020) and other dicot species from the PLAZA database (Proost et al., 2015).

### Sequence availability

Sequencing data for the *RPB1* gene containing regions of Est-1 and Uod-1 are available from GenBank under the entries ON529272 and ON529273 respectively.

## SUPPLEMENTARY FIGURE LEGENDS

**Figure S1. Development of clubroot galls and accumulation of *P. brassicae* DNA over course of infection.** A) Morphology of clubroot galls developing in Arabidopsis accession Col-0, scale bar represents 10 mm. B) Quantification of pathogen DNA titre relative to host DNA by qPCR (*Pb18S/AtSK11*). Four biological replicates each consisting of 3 plants were harvested at each time-point, at 7 dpi *P. brassicae* DNA was below the limit of detection in 2 replicates.

**Figure S2. Clubroot symptoms in Arabidopsis accessions Pro-0 and Var-2-6.** A) Rosette symptoms in *P. brassicae* infected Pro-0 19 dpi. B) Clubroot gall development in Pro-0 19 dpi, squares on the background grid have 10 mm length. C) Rosette symptoms in *P. brassicae* infected Var-2-6 19 dpi. D) Clubroot gall development in Var-2-6 19 dpi, squares on the background grid have 10 mm length.

**Figure S3. Clubroot resistance associated SNP adjacent to *WIP2*.** A) Overview of polymorphisms in the sequence 1225 bp up-stream of the gene WIP2, comparing Col-0 with resistant accessions HR-10, Pu2-23, Tsu-0 and Uod-1. B & C) Expression of *WIP2* and *AT4G05071* 7 dpi relative to *AT1G76030* and *AT3G48140*. Each point represents one biological replicate (n = 4) of 10 plants, horizontal bars indicate the mean.

**Figure S4. Sequence polymorphisms in *RAC1*.** A) Alignment of the predicted protein sequence for the N-terminal region of RAC1 in various clubroot resistant (R) or susceptible (S) accessions. B) Expression of *RAC1* 7 dpi relative to *AT1G76030* and *AT3G48140*. Different letters indicate statistically significant differences (BH adjusted p < 0.05). Each point represents one biological replicate (n = 4) of 10 plants, horizontal bars indicate the mean.

**Figure S5. Deletion of *RAC1* does not affect clubroot resistance in Est-1 or Uod-1.** A) Gall and rosette symptoms of *P. brassicae* infection 19 dpi for *rac1* knock-out mutants. White bars = 2 cm. B) Relative pathogen DNA titre (*Pb18S/AtSK11*) 19 dpi, points indicate biological replicates of 3 plants, horizontal lines indicate the means, different letters indicate statistically significant differences (Tukey, BH adjusted p < 0.05), n = 6. C) DI score for the symptoms of galls in B, the percentage of plants assigned to each symptom class is shown on each segment, n = 18.

**Figure S6. RPB1 protein sequence is conserved across clubroot susceptible and resistant accessions.** Alignment of predicted amino acid sequence for *RPB1* genes from clubroot resistant (Tsu-0, Est-1, Uod-1) and susceptible (An-1, C24, Kyoto, Cvi-0) accessions, RLD-1 was not tested.

**Figure S7. Restriction of clubroot gall development in Est-1 and Uod-1 is dependent on *RPB1*.** Representative mock (A) and *P. brassicae* (B) inoculated hypocotyls and roots from wildtype and *rpb1* knock-out lines 25 dpi. Scale bar = 2 mm.

**Figure S8. Deletion of *RPB1* does not affect growth or development.** Histological characterization of the hypocotyls of mock inoculated wildtype accessions and *rpb1* mutants 25 dpi for comparison with Figure 4. Micrographs in the left column focus on the uppermost central part of the hypocotyl, the central column focusses on the epidermal cells. In the right column the region of the interphase between the hypocotyl and the root is presented. X: Xylem cells. The scale bar in the inset overview picture corresponds to 1000 μm, the scale bar in the detailed pictures corresponds to 50 μm.

**Figure S9. *RPB1* is required for lignification of host cell walls in response to *P. brassicae*.** Detailed visualisation of pathogen structures surrounded by lignified cells in the (A) Uod-1 and (B) Est-1 wildtype genotypes inoculated with *P. brassicae* 25 dpi. L: Lignified tissue, Pb: *P. brassicae* cells. The scale bar corresponds to 50 μm.

**Figure S10. RPB1 is predicted to contain membrane spanning domains.** Prediction of membrane spanning domains (purple) in the sequence of RPB1 made by Phobius.

**Figure S11. RPB1 orthologues are present in Brassicaceae species.** Identification of putative orthologues of *RPB1* and *RPB1-like* genes in Brassicacea species by OrthoFinder. All Arabidopsis *RPB1* related sequences were grouped in one orthogroup.

## SUPPLEMENTAL TABLES

**Table S1. Phenotype data for Arabidopsis accessions infected with *P. brassicae*.**

**Table S2. Mutations generated in *RAC1*.**

**Table S3. Primers used in this study.**

## AKNOWLEDGEMENTS

This work was supported by funding from the National Science Centre, Poland - OPUS UMO-2015/19/B/NZ3/01489 and PRELUDIUM UMO-2019/33/N/NZ9/01048.

## Notes

### Competing Interest Statement

The authors have declared no competing interest.

## REFERENCES

Alix K, Lariagon C, Delourme R, Manzanares-Dauleux MJ (2007) Exploiting natural genetic diversity and mutant resources of Arabidopsis thaliana to study the *A*. thaliana Plasmodiophora brassicae interaction. Plant Breeding 126: 218–221

Alonso-Blanco C, Andrade J, Becker C, Bemm F, Bergelson J, Borgwardt KM, Cao J, Chae E, Dezwaan TM, Ding W, et al (2016) 1,135 Genomes Reveal the Global Pattern of Polymorphism in Arabidopsis thaliana. Cell 166: 481–491

Arbeiter A, Fähling M, Graf H, Sacristán MD, Siemens J (2002) Resistance of Arabidopsis thaliana to the obligate biotrophic parasite Plasmodiophora brassicae. Plant Protect Sci 38: 519–522

Bates D, Mächler M, Bolker B, Walker S (2015) Fitting Linear Mixed-Effects Models Using lme4. J Stat Soft. doi: 10.18637/jss.v067.i01

Benjamini Y, Hochberg Y (1995) Controlling the False Discovery Rate: A Practical and Powerful Approach to Multiple Testing. Journal of the Royal Statistical Society: Series B (Methodological) 57: 289–300

Bieluszewski T, Sura W, Dziegielewski W, Bieluszewska A, Lachance C, Kabza M, Szymanska-Lejman M, Abram M, Wlodzimierz P, De Winne N, et al (2022) NuA4 and H2A.Z control environmental responses and autotrophic growth in Arabidopsis. Nat Commun 13: 127

Bland JM, Altman DG (1995) Statistics notes: Multiple significance tests: the Bonferroni method. BMJ 310: 170–170

Borhan MH, Holub EB, Beynon JL, Rozwadowski K, Rimmer SR (2004) The Arabidopsis TIR-NB-LRR Gene *RAC1* Confers Resistance to *Albugo candida* (White Rust) and Is Dependent on *EDS1* but not *PAD4*. MPMI 17: 711–719

Breitenbach HH, Wenig M, Wittek F, Jordá L, Maldonado-Alconada AM, Sarioglu H, Colby T, Knappe C, Bichlmeier M, Pabst E, et al (2014) Contrasting Roles of the Apoplastic Aspartyl Protease APOPLASTIC, *ENHANCED DISEASE SUSCEPTIBILITY1*-DEPENDENT1 and LEGUME LECTIN-LIKE PROTEIN1 in Arabidopsis Systemic Acquired Resistance,. Plant Physiology 165: 791–809

Buczacki ST, Toxopeus H, Mattusch P, Johnston TD, Dixon GR, Hobolth LA (1975) Study of physiologic specialization in Plasmodiophora brassicae: Proposals for attempted rationalization through an international approach. Transactions of the British Mycological Society 65: 295–303

Bulman S, Richter F, Marschollek S, Benade F, Jülke S, Ludwig-Müller J (2019) *Arabidopsis thaliana* expressing *Pb BSMT*, a gene encoding a SABATH-type methyltransferase from the plant pathogenic protist *Plasmodiophora brassicae*, show leaf chlorosis and altered host susceptibility. Plant Biol J 21: 120–130

Chen H, Wang T, He X, Cai X, Lin R, Liang J, Wu J, King G, Wang X (2022) BRAD V3.0: an upgraded Brassicaceae database. Nucleic Acids Research 50: D1432–D1441

Choi K, Reinhard C, Serra H, Ziolkowski PA, Underwood CJ, Zhao X, Hardcastle TJ, Yelina NE, Griffin C, Jackson M, et al (2016) Recombination Rate Heterogeneity within Arabidopsis Disease Resistance Genes. PLoS Genet 12: e1006179

Clough SJ, Bent AF (1998) Floral dip: a simplified method forAgrobacterium-mediated transformation ofArabidopsis thaliana: Floral dip transformation of Arabidopsis. The Plant Journal 16: 735–743

Concordet J-P, Haeussler M (2018) CRISPOR: intuitive guide selection for CRISPR/Cas9 genome editing experiments and screens. Nucleic Acids Research 46: W242–W245

Corwin JA, Copeland D, Feusier J, Subedy A, Eshbaugh R, Palmer C, Maloof J, Kliebenstein DJ (2016) The Quantitative Basis of the Arabidopsis Innate Immune System to Endemic Pathogens Depends on Pathogen Genetics. PLoS Genet 12: e1005789

Diederichsen E, Beckmann J, Schondelmeier J, Dreyer F (2006) GENETICS OF CLUBROOT RESISTANCE IN BRASSICA NAPUS ‘MENDEL’. Acta Hortic 307–312

Diederichsen E, Frauen M, Linders EGA, Hatakeyama K, Hirai M (2009) Status and Perspectives of Clubroot Resistance Breeding in Crucifer Crops. J Plant Growth Regul 28: 265–281

Emms DM, Kelly S (2019) OrthoFinder: phylogenetic orthology inference for comparative genomics. Genome Biol 20: 238

Fredua-Agyeman R, Rahman H (2016) Mapping of the clubroot disease resistance in spring Brassica napus canola introgressed from European winter canola cv. ‘Mendel.’ Euphytica 211: 201–213

Fuchs H and Sacristan MD (1996) Identification of a Gene in *Arabidopsis thaliana* Controlling Resistance to Clubroot (*Plasmodiophora brassicae*) and Characterization of the Resistance Response. MPMI 9: 091

Galindo-González L, Manolii V, Hwang S-F, Strelkov SE (2020) Response of Brassica napus to Plasmodiophora brassicae Involves Salicylic Acid-Mediated Immunity: An RNA-Seq-Based Study. Front Plant Sci 11: 1025

Gravot A, Grillet L, Wagner G, Jubault M, Lariagon C, Baron C, Deleu C, Delourme R, Bouchereau A, Manzanares-Dauleux MJ (2011) Genetic and physiological analysis of the relationship between partial resistance to clubroot and tolerance to trehalose in *Arabidopsis thaliana*. New Phytologist 191: 1083–1094

Grimm DG, Roqueiro D, Salomé PA, Kleeberger S, Greshake B, Zhu W, Liu C, Lippert C, Stegle O, Schölkopf B, et al (2017) easyGWAS: A Cloud-Based Platform for Comparing the Results of Genome-Wide Association Studies. Plant Cell 29: 5–19

Hatakeyama K, Suwabe K, Tomita RN, Kato T, Nunome T, Fukuoka H, Matsumoto S (2013) Identification and Characterization of Crr1a, a Gene for Resistance to Clubroot Disease (Plasmodiophora brassicae Woronin) in Brassica rapa L. PLoS ONE 8: e54745

Horton MW, Hancock AM, Huang YS, Toomajian C, Atwell S, Auton A, Muliyati NW, Platt A, Sperone FG, Vilhjálmsson BJ, et al (2012) Genome-wide patterns of genetic variation in worldwide Arabidopsis thaliana accessions from the RegMap panel. Nat Genet 44: 212–216

Ji Z, Guo W, Chen X, Wang C, Zhao K (2022) Plant Executor Genes. IJMS 23: 1524

Jiao W-B, Schneeberger K (2020) Chromosome-level assemblies of multiple Arabidopsis genomes reveal hotspots of rearrangements with altered evolutionary dynamics. Nat Commun 11: 989

Jubault M, Lariagon C, Simon M, Delourme R, Manzanares-Dauleux MJ (2008) Identification of quantitative trait loci controlling partial clubroot resistance in new mapping populations of Arabidopsis thaliana. Theor Appl Genet 117: 191–202

Käll L, Krogh A, Sonnhammer ELL (2004) A Combined Transmembrane Topology and Signal Peptide Prediction Method. Journal of Molecular Biology 338: 1027–1036

Lemarié S, Robert-Seilaniantz A, Lariagon C, Lemoine J, Marnet N, Jubault M, Manzanares-Dauleux MJ, Gravot A (2015a) Both the Jasmonic Acid and the Salicylic Acid Pathways Contribute to Resistance to the Biotrophic Clubroot Agent *Plasmodiophora brassicae* in Arabidopsis. Plant Cell Physiol pcv 127

Lemarié S, Robert-Seilaniantz A, Lariagon C, Lemoine J, Marnet N, Levrel A, Jubault M, Manzanares-Dauleux MJ, Gravot A (2015b) Camalexin contributes to the partial resistance of Arabidopsis thaliana to the biotrophic soilborne protist Plasmodiophora brassicae. Front Plant Sci. doi: 10.3389/fpls.2015.00539

Liu L, Qin L, Cheng X, Zhang Y, Xu L, Liu F, Tong C, Huang J, Liu S, Wei Y (2020a) Comparing the Infection Biology of Plasmodiophora brassicae in Clubroot Susceptible and Resistant Hosts and Non-hosts. Front Microbiol 11: 507036

Liu L, Qin L, Zhou Z, Hendriks WGHM, Liu S, Wei Y (2020b) Refining the Life Cycle of *Plasmodiophora brassicae*. Phytopathology 110: 1704–1712

Ludwig-Müller J, Auer S, Jülke S, Marschollek S (2017) Manipulation of Auxin and Cytokinin Balance During the Plasmodiophora brassicae–Arabidopsis thaliana Interaction. In T Dandekar, M Naseem, eds, Auxins and Cytokinins in Plant Biology. Springer New York, New York, NY, pp 41–60

Malinowski R, Truman W, Blicharz S (2019) Genius Architect or Clever Thief—How *Plasmodiophora brassicae* Reprograms Host Development to Establish a Pathogen-Oriented Physiological Sink. MPMI 32: 1259–1266

Mehraj H, Akter A, Miyaji N, Miyazaki J, Shea DJ, Fujimoto R, Doullah MdA (2020) Genetics of Clubroot and Fusarium Wilt Disease Resistance in Brassica Vegetables: The Application of Marker Assisted Breeding for Disease Resistance. Plants 9: 726

Nafisi M, Goregaoker S, Botanga CJ, Glawischnig E, Olsen CE, Halkier BA, Glazebrook J (2007) *Arabidopsis* Cytochrome P450 Monooxygenase 71A13 Catalyzes the Conversion of Indole-3-Acetaldoxime in Camalexin Synthesis. The Plant Cell 19: 2039–2052

Norkunas K, Harding R, Dale J, Dugdale B (2018) Improving agroinfiltration-based transient gene expression in Nicotiana benthamiana. Plant Methods 14: 71

Olszak M, Truman W, Stefanowicz K, Sliwinska E, Ito M, Walerowski P, Rolfe S, Malinowski R (2019) Transcriptional profiling identifies critical steps of cell cycle reprogramming necessary for *Plasmodiophora brassicae*-driven gall formation in Arabidopsis. Plant J 97: 715–729

Peng G, Lahlali R, Hwang S-F, Pageau D, Hynes RK, McDonald MR, Gossen BD, Strelkov SE (2014) Crop rotation, cultivar resistance, and fungicides/biofungicides for managing clubroot (*Plasmodiophora brassicae*) on canola. Canadian Journal of Plant Pathology 36: 99–112

Peterson R A (2021) Finding Optimal Normalizing Transformations via bestNormalize. The R Journal 13: 310

Pina C, Pinto F, Feijó JA, Becker JD (2005) Gene Family Analysis of the Arabidopsis Pollen Transcriptome Reveals Biological Implications for Cell Growth, Division Control, and Gene Expression Regulation. Plant Physiology 138: 744–756

Proost S, Van Bel M, Vaneechoutte D, Van de Peer Y, Inzé D, Mueller-Roeber B, Vandepoele K (2015) PLAZA 3.0: an access point for plant comparative genomics. Nucleic Acids Research 43: D974–D981

Ramzi N, Kaczmarek J, Jedryczka M (2018) Identification of clubroot resistance sources from world gene bank accessions. IOBC/WPRS Bulletin 136: 144–147

Řičařová V, Kaczmarek J, Strelkov SE, Kazda J, Lueders W, Rysanek P, Manolii V, Jedryczka M (2016) Pathotypes of Plasmodiophora brassicae causing damage to oilseed rape in the Czech Republic and Poland. Eur J Plant Pathol 145: 559–572

Rolfe SA, Strelkov SE, Links MG, Clarke WE, Robinson SJ, Djavaheri M, Malinowski R, Haddadi P, Kagale S, Parkin IAP, et al (2016) The compact genome of the plant pathogen Plasmodiophora brassicae is adapted to intracellular interactions with host Brassica spp. BMC Genomics 17: 272

Rousseau-Gueutin M, Belser C, Da Silva C, Richard G, Istace B, Cruaud C, Falentin C, Boideau F, Boutte J, Delourme R, et al (2020) Long-read assembly of the *Brassica napus* reference genome Darmor-bzh. GigaScience 9: giaa137

Schneider CA, Rasband WS, Eliceiri KW (2012) NIH Image to ImageJ: 25 years of image analysis. Nat Methods 9: 671–675

Seren Ü, Vilhjálmsson BJ, Horton MW, Meng D, Forai P, Huang YS, Long Q, Segura V, Nordborg M (2013) GWAPP: A Web Application for Genome-Wide Association Mapping in Arabidopsis. The Plant Cell 24: 4793–4805

Serra H, Choi K, Zhao X, Blackwell AR, Kim J, Henderson IR (2018) Interhomolog polymorphism shapes meiotic crossover within the Arabidopsis RAC1 and RPP13 disease resistance genes. PLoS Genet 14: e1007843

Sharma K, Gossen BD, Greenshields D, Selvaraj G, Strelkov SE, McDonald MR (2013) Reaction of Lines of the Rapid Cycling Brassica Collection and *Arabidopsis thaliana* to Four Pathotypes of *Plasmodiophora brassicae*. Plant Disease 97: 720–727

Some A, Manzanares MJ, Laurens F, Baron F, Thomas G, Rouxel F (1996) Variation for virulence on *Brassica napus* L. amongst *Plasmodiophora brassicae* collections from France and derived single-spore isolates. Plant Pathology 45: 432–439

Stefanowicz K, Szymanska-Chargot M, Truman W, Walerowski P, Olszak M, Augustyniak A, Kosmala A, Zdunek A, Malinowski R (2021) Plasmodiophora brassicae-Triggered Cell Enlargement and Loss of Cellular Integrity in Root Systems Are Mediated by Pectin Demethylation. Front Plant Sci 12: 711838

Strelkov SE, Hwang S-F, Manolii VP, Cao T, Feindel D (2016) Emergence of new virulence phenotypes of Plasmodiophora brassicae on canola (Brassica napus) in Alberta, Canada. Eur J Plant Pathol 145: 517–529

Struck C, Rüsch S, Strehlow B (2022) Control Strategies of Clubroot Disease Caused by Plasmodiophora brassicae. Microorganisms 10: 620

Ueno H, Matsumoto E, Aruga D, Kitagawa S, Matsumura H, Hayashida N (2012) Molecular characterization of the CRa gene conferring clubroot resistance in Brassica rapa. Plant Mol Biol 80: 621–629

Walerowski P, Gündel A, Yahaya N, Truman W, Sobczak M, Olszak M, Rolfe S, Borisjuk L, Malinowski R (2018) Clubroot Disease Stimulates Early Steps of Phloem Differentiation and Recruits SWEET Sucrose Transporters within Developing Galls. Plant Cell 30: 3058–3073

Wallenhammar A-C. (1996) Prevalence of *Plasmodiophora brassicae* in a spring oilseed rape growing area in central Sweden and factors influencing soil infestation levels. Plant Pathology 45: 710–719

Zamani-Noor N, Brand S, Söchting H-P (2022) Effect of Pathogen Virulence on Pathogenicity, Host Range, and Reproduction of *Plasmodiophora brassicae*, the Causal Agent of Clubroot Disease. Plant Disease 106: 57–64

Zamani-Noor N, Hornbacher J, Comel CJ, Papenbrock J (2021) Variation of Glucosinolate Contents in Clubroot-Resistant and - Susceptible Brassica napus Cultivars in Response to Virulence of Plasmodiophora brassicae. Pathogens 10: 563

Zhou Q, Galindo-González L, Manolii V, Hwang S-F, Strelkov SE (2020) Comparative Transcriptome Analysis of Rutabaga (Brassica napus) Cultivars Indicates Activation of Salicylic Acid and Ethylene-Mediated Defenses in Response to Plasmodiophora brassicae. IJMS 21: 8381

Zimmermann P, Hirsch-Hoffmann M, Hennig L, Gruissem W (2004) GENEVESTIGATOR. Arabidopsis Microarray Database and Analysis Toolbox. Plant Physiology 136: 2621–2632

