## Supplemental Figure 1 for "Natural variation in Arabidopsis responses to *Plasmodiophora brassicae* reveals an essential role for RPB1"

A

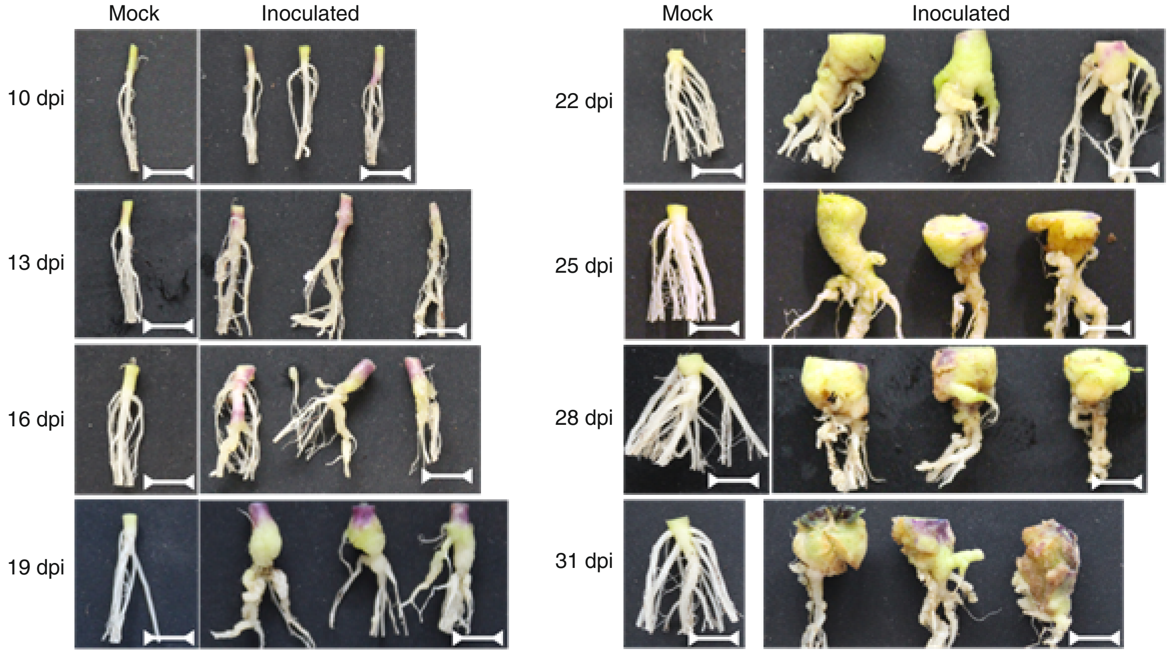


B

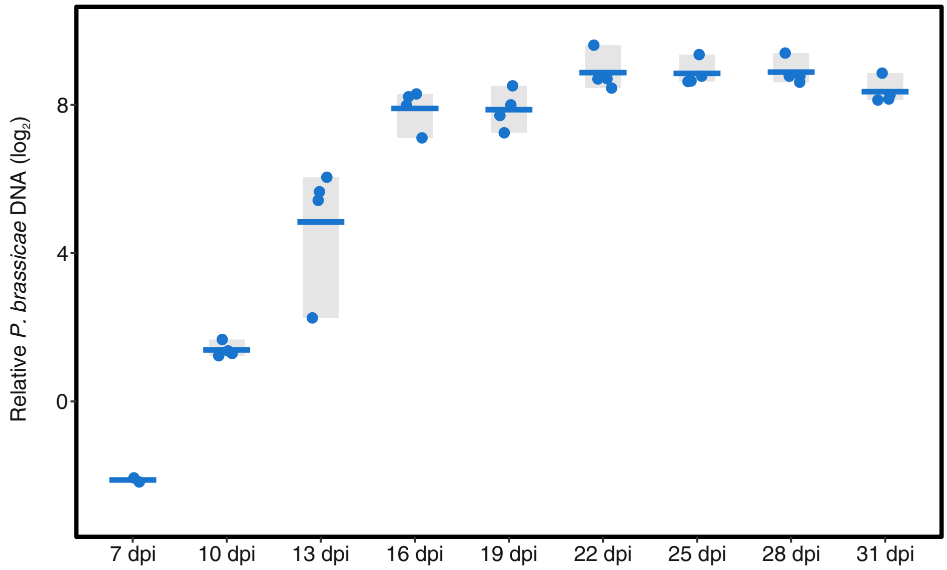


**Figure S1. Development of clubroot galls and accumulation of *P. brassicae* DNA over course of infection.**

A) Morphology of clubroot galls developing in Arabidopsis accession Col-0, scale bar represents 10 mm. B) Quantification of pathogen DNA titre relative to host DNA by qPCR (*Pb18S/AtSK11*). Four biological replicates each consisting of 3 plants were harvested at each time-point, at 7 dpi *P. brassicae* DNA was below the limit of detection in 2 replicates.
