## Supplemental Figure 2 for "Natural variation in Arabidopsis responses to *Plasmodiophora brassicae* reveals an essential role for RPB1"

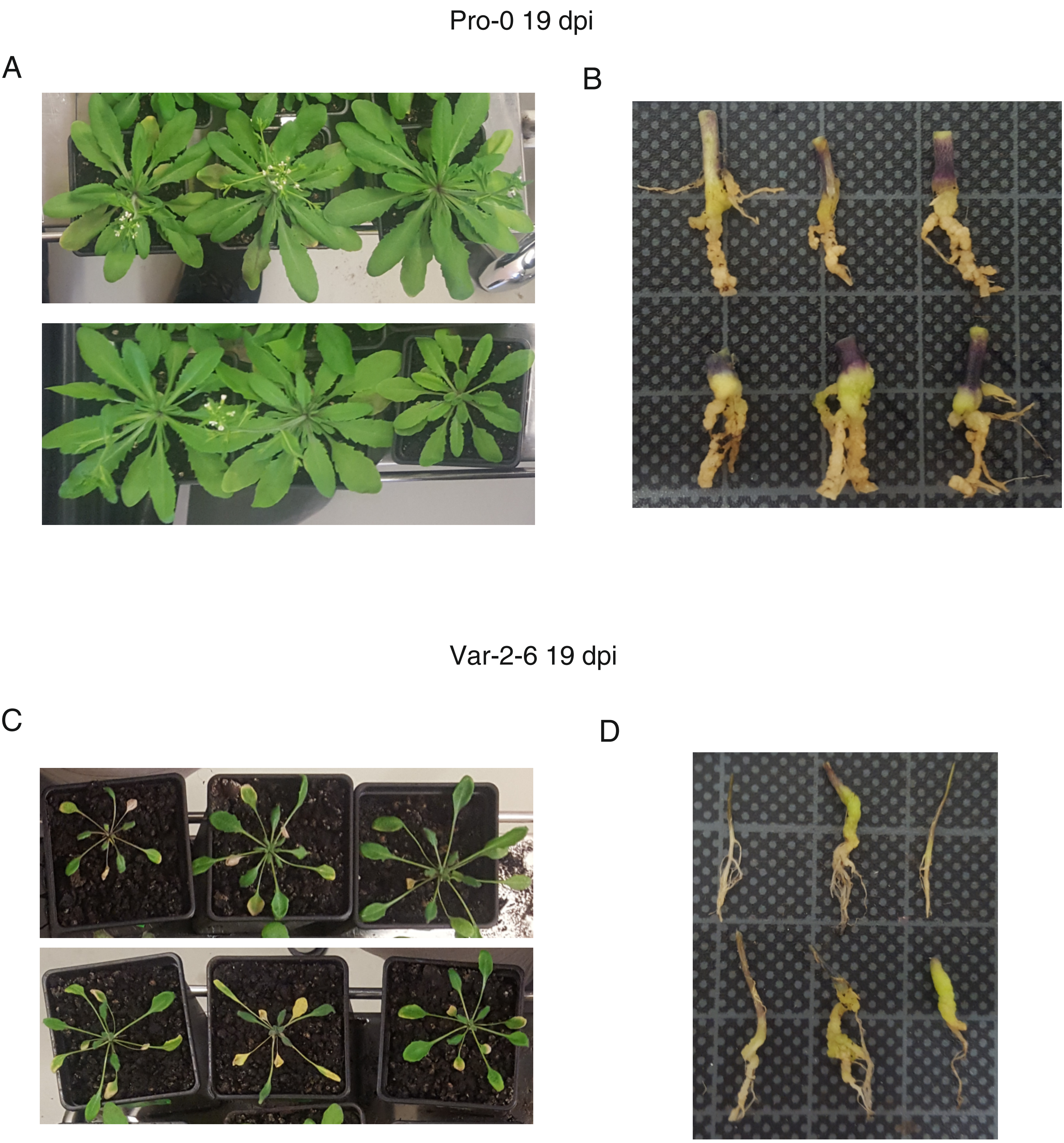


**Figure S2. Clubroot symptoms in Arabidopsis accessions Pro-0 and Var-2-6.**
A) Rosette symptoms in *P. brassicae* infected Pro-0 19 dpi. B) Clubroot gall development in Pro-0 19 dpi, squares on the background grid have 10 mm length. C) Rosette symptoms in *P. brassicae* infected Var-2-6 19 dpi. D) Clubroot gall development in Var-2-6 19 dpi, squares on the background grid have 10 mm length.
