## Supplemental Figure 3 for "Natural variation in Arabidopsis responses to *Plasmodiophora brassicae* reveals an essential role for RPB1"

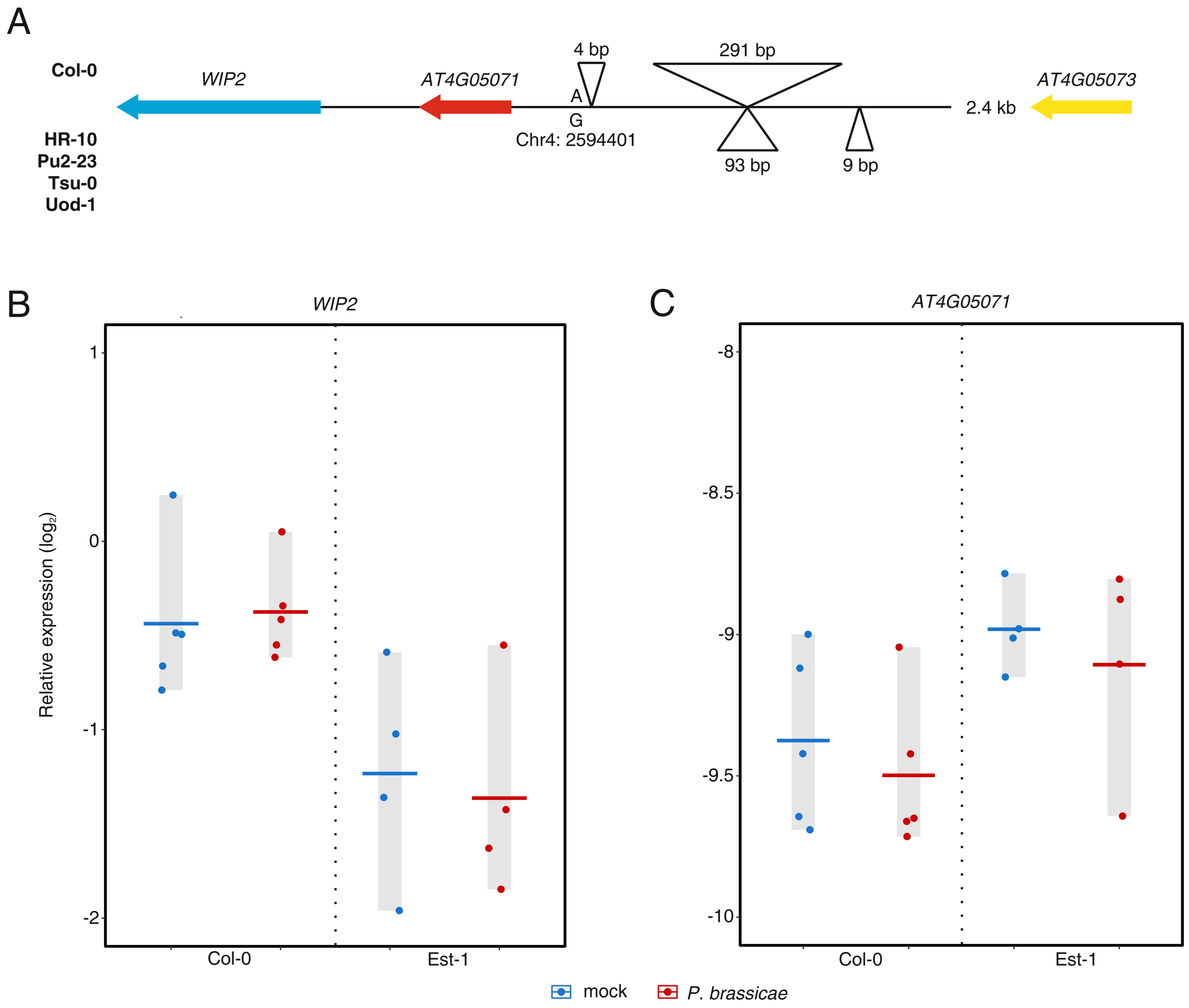


**Figure S3. Clubroot resistance associated SNP adjacent to *WIP2*.**
A) Overview of polymorphisms in the sequence 1225 bp up-stream of the gene WIP2, comparing Col-0 with resistant accessions HR-10, Pu2-23, Tsu-0 and Uod-1. B & C) Expression of *WIP2* and *AT4G05071* 7 dpi relative to *AT1G76030* and *AT3G48140*. Each point represents one biological replicate (n = 4) of 10 plants, horizontal bars indicate the mean.
