## Supplemental Figure 4 for "Natural variation in Arabidopsis responses to *Plasmodiophora brassicae* reveals an essential role for RPB1"

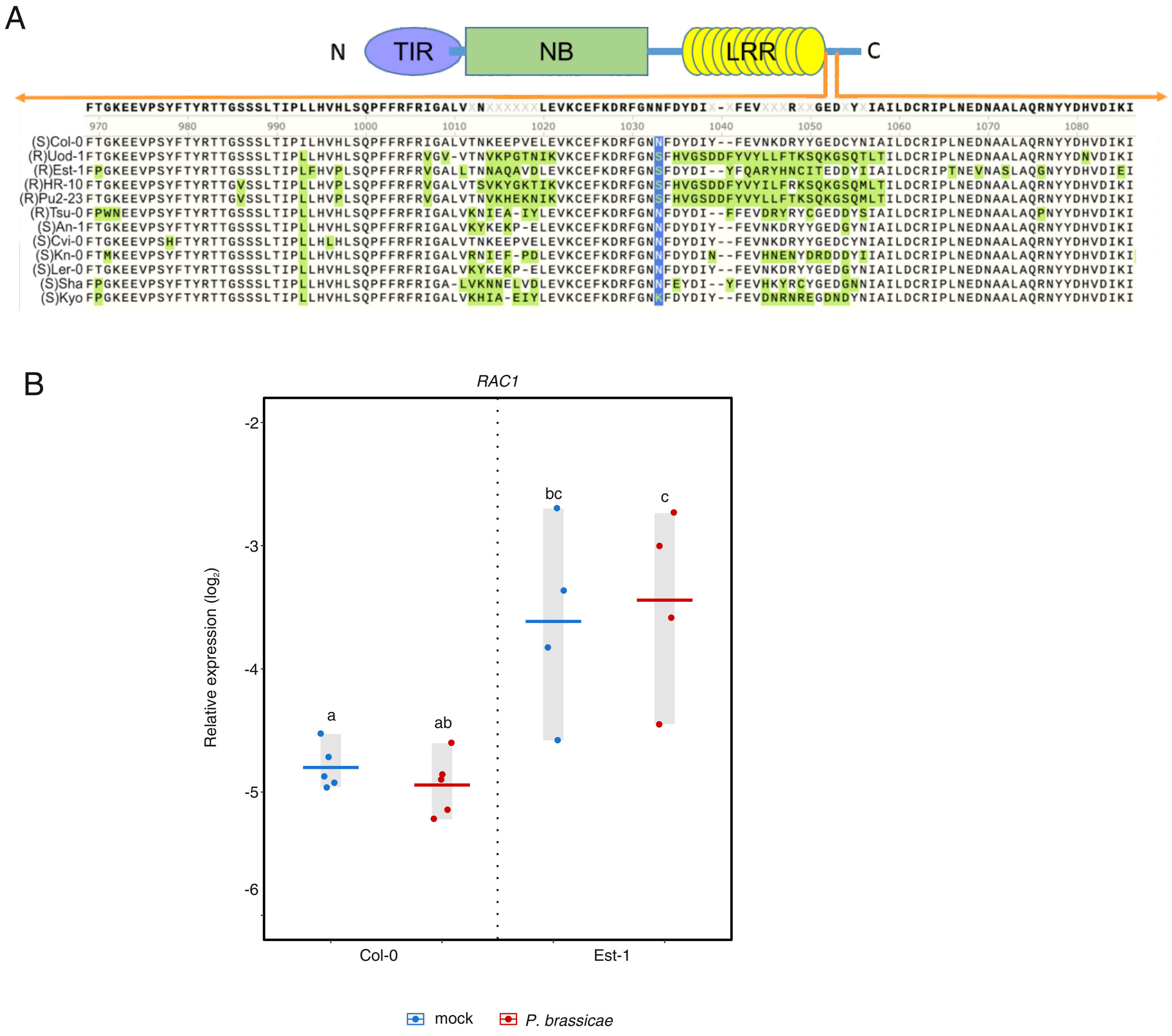


**Figure S4. Sequence polymorphisms in *RAC1*.**

A) Alignment of the predicted protein sequence for the N-terminal region of RAC1 in various clubroot resistant (R) or susceptible (S) accessions. B) Expression of *RAC1* 7 dpi relative to *AT1G76030* and *AT3G48140*. Different letters indicate statistically significant differences (BH adjusted $p < 0.05$). Each point represents one biological replicate ($n = 4$) of 10 plants, horizontal bars indicate the mean.
