## Supplemental Figure 5 for "Natural variation in Arabidopsis responses to *Plasmodiophora brassicae* reveals an essential role for RPB1"

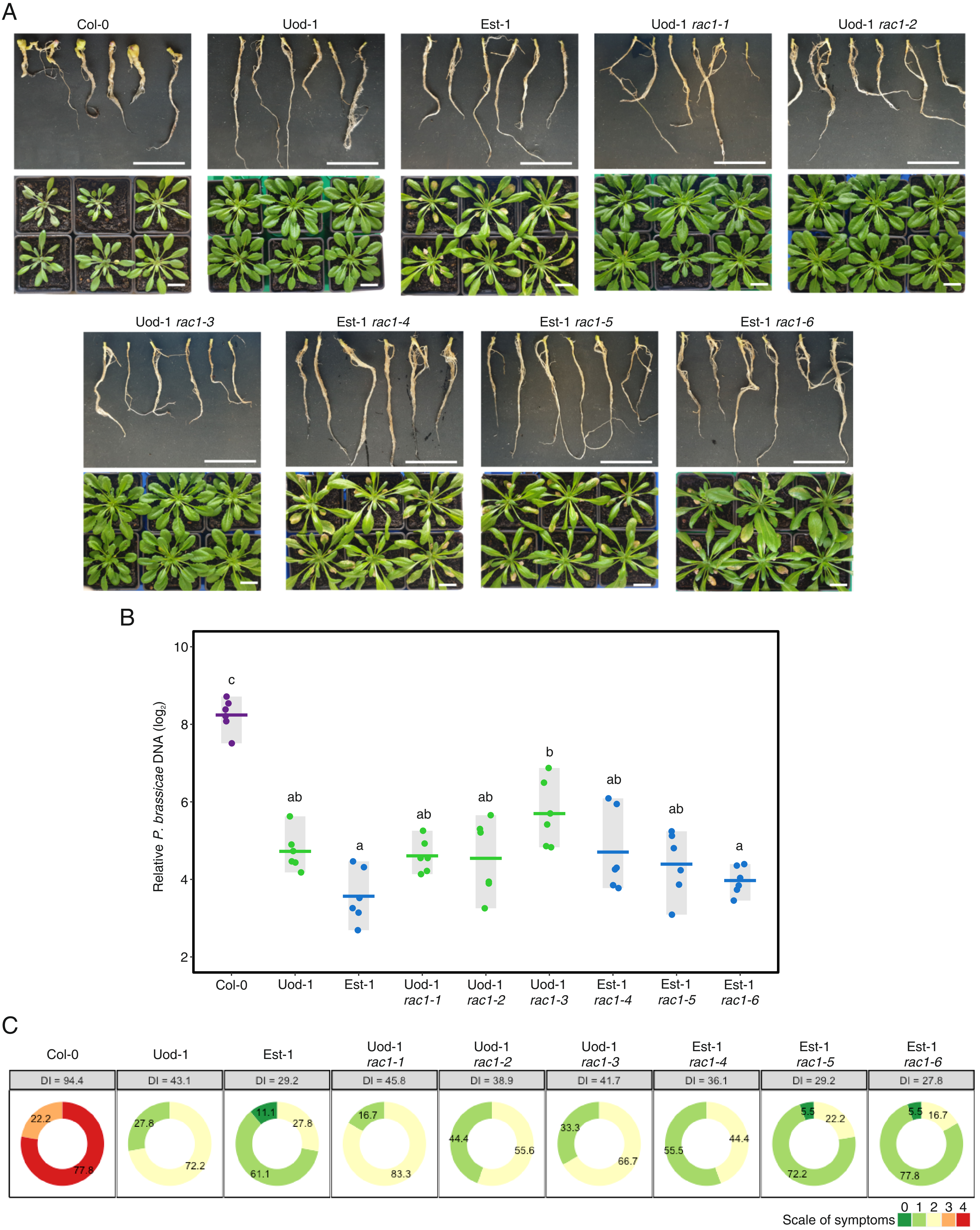


**Figure S5. Deletion of *RAC1* does not affect clubroot resistance in Est-1 or Uod-1.**
A) Gall and rosette symptoms of *P. brassicae* infection 19 dpi for *rac1* knock-out mutants. White bars = 2 cm. B) Relative pathogen DNA titre (*Pb18S/AtSK11*) 19 dpi, points indicate biological replicates of 3 plants, horizontal lines indicate the means, different letters indicate statistically significant differences (Tukey, BH adjusted $p < 0.05$), n = 6. C) DI score for the symptoms of galls in B, the percentage of plants assigned to each symptom class is shown on each segment, n = 18.
