## Supplemental Figure 6 for "Natural variation in Arabidopsis responses to *Plasmodiophora brassicae* reveals an essential role for RPB1"

```
Tsu0_RPB1a          METVSAVNQTLPISGGEPVKFTTYSAAVHKVLVMVNAGILGLLQLVSQQSSVLETHKAAF     60
Tsu0_RPB1b          METVSAVNQTLPISGGEPVKFTTYSAAVHKVLVMVNAGILGLLQLVSQQSSVLETHKAAF     60
RLD-1_RPB1          METVSAVNQTLPISGGEPVKFTTYSAAVHKVLVMVNAGILGLLQLVSQQSSVLETHKAAF     60
Est-1_RPB1          METVSAVNQTLPISGGEPVKFTTYSAAVHKVLVMVNAGILGLLQLVSQQSSVLETHKAAF     60
Uod-1_RPB1          METVSAVNQTLPISGGEPVKFTTYSAAVHKVLVMVNAGILGLLQLVSQQSSVLETHKAAF     60
ATAN-1G43570.1      METVSAVNQTLPISGGEPVKFTTYSAAVHKVLVMVNAGILGLLQLVSQQSSVLETHKAAF     60
ATC24-1G43720.1     METVSAVNQTLPISGGEPVKFTTYSAAVHKVLVMVNAGILGLLQLVSQQSSVLETHKAAF     60
ATKYO-1G43930.1     METVSAVNQTLPISGGEPVKFTTYSAAVHKVLVMVNAGILGLLQLVSQQSSVLETHKAAF     60
ATCVI-1G44000.1     METVSAVNQTLPISGGEPVKFTTYSAAVHKVLVMVNAGILGLLQLVSQQSSVLETHKAAF     60
                    ************************************************************

Tsu0_RPB1a          LCFCVFILFYAVLRVREAMDVRLQPGLVPRLIGHGSHLFGGLAALVLVSVVSTAFSIVLF     120
Tsu0_RPB1b          LCFCVFILFYAVLRVREAMDVRLQPGLVPRLIGHGSHLFGGLAALVLVSVVSTAFSIVLF     120
RLD-1_RPB1          LCFCVFILFYAVLRVREAMDVRLQPGLVPRLIGHGSHLFGGLAALVLVSVVSTAFSIVLF     120
Est-1_RPB1          LCFCVFILFYAVLRVREAMDVRLQPGLVPRLIGHGSHLFGGLAALVLVSVVSTAFSIVLF     120
Uod-1_RPB1          LCFCVFILFYAVLRVREAMDVRLQPGLVPRLIGHGSHLFGGLAALVLVSVVSTAFSIVLF     120
ATAN-1G43570.1      LCFCVFILFYAVLRVREAMDVRLQPGLVPRLIGHGSHLFGGLAALVLVSVVSTAFSIVLF     120
ATC24-1G43720.1     LCFCVFILFYAVLRVREAMDVRLQPGLVPRLIGHGSHLFGGLAALVLVSVVSTAFSIVLF     120
ATKYO-1G43930.1     LCFCVFILFYAVLRVREAMDVRLQPGLVPRLIGHGSHLFGGLAALVLVSVVSTAFSIVLF     120
ATCVI-1G44000.1     LCFCVFILFYAVLRVREAMDVRLQPGLVPRLIGHGSHLFGGLAALVLVSVVSTAFSIVLF     120
                    ************************************************************

Tsu0_RPB1a          LLWFIWLSAVVYLETNKPSACPPQLPPV        148
Tsu0_RPB1b          LLWFIWLSAVVYLETNKPSACPPQLPPV        148
RLD-1_RPB1          LLWFIWLSAVVYLETNKPSACPPQLPPV        148
Est-1_RPB1          LLWFIWLSAVVYLETNKPSACPPQLPPV        148
Uod-1_RPB1          LLWFIWLSAVVYLETNKPSACPPQLPPV        148
ATAN-1G43570.1      LLWFIWLSAVVYLETNKPSACPPQLPPV        148
ATC24-1G43720.1     LLWFIWLSAVVYLETNKPSACPPQLPPV        148
ATKYO-1G43930.1     LLWFIWLSAVVYLETNKPSACPPQLPPV        148
ATCVI-1G44000.1     LLWFIWLSAVVYMETNKPSACPPQLPPV        148
                    ************:***************
```

**Figure S6. RPB1 protein sequence is conserved across clubroot susceptible and resistant accessions.**

Alignment of predicted amino acid sequence for *RPB1* genes from clubroot resistant (Tsu-0, Est-1, Uod-1) and susceptible (An-1, C24, Kyoto, Cvi-0) accessions, RLD-1 was not tested.
