## Supplemental Figure 7 for "Natural variation in Arabidopsis responses to *Plasmodiophora brassicae* reveals an essential role for RPB1"

A

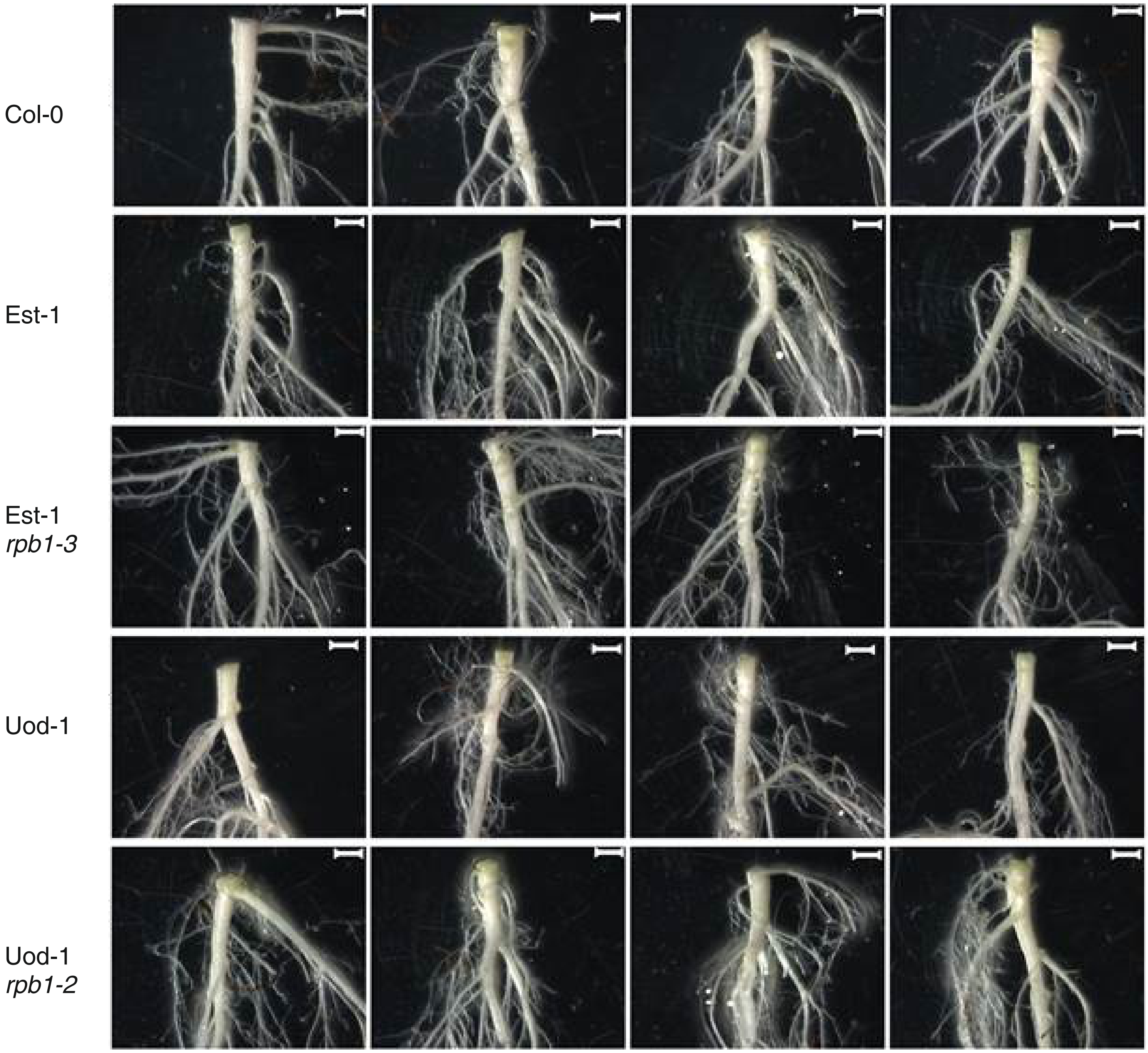

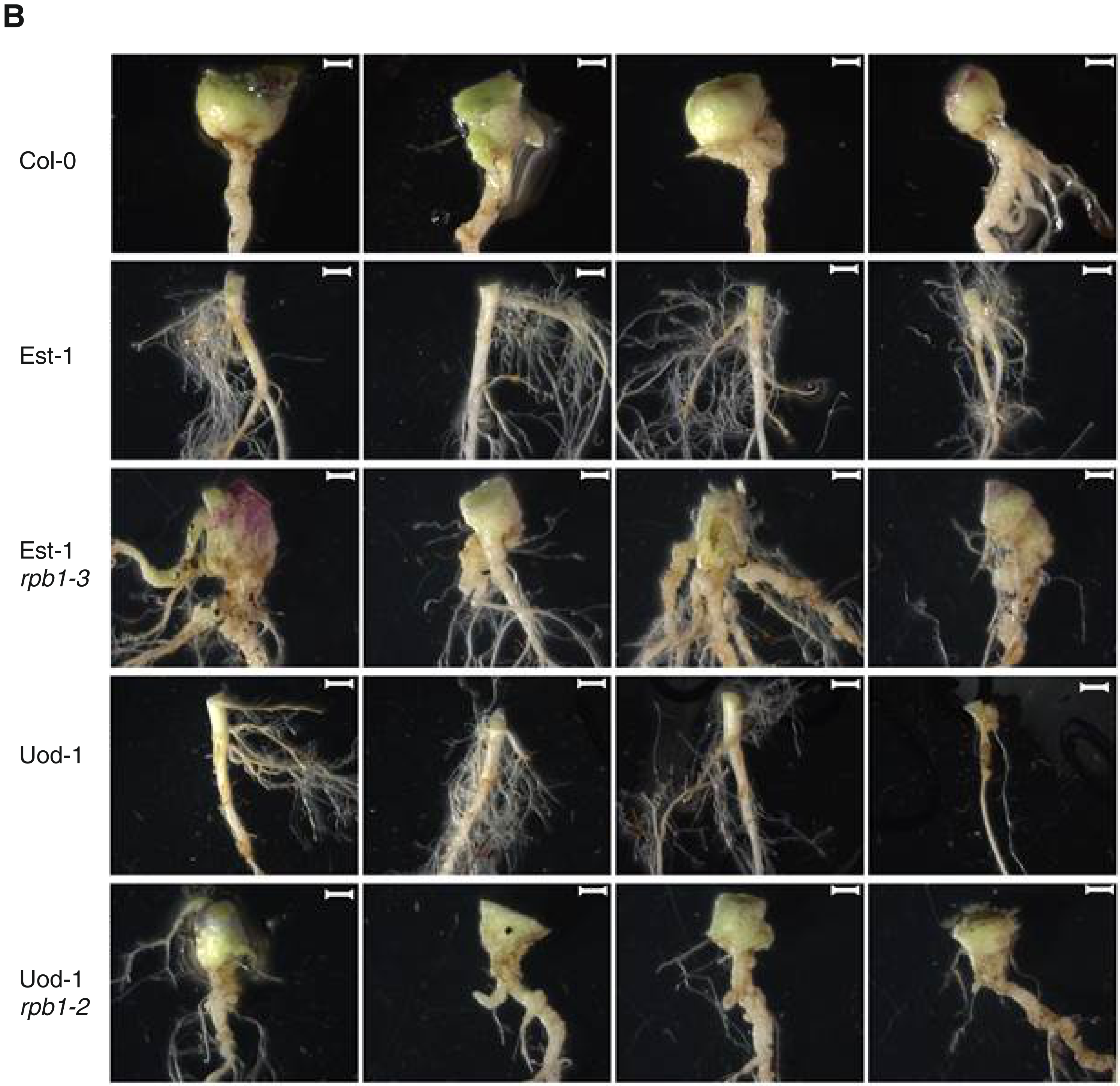


**Figure S7. Restriction of clubroot gall development in Est-1 and Uod-1 is dependent on *RPB1*.**
Representative mock (A) and *P. brassicae* (B) inoculated hypocotyls and roots from wildtype and *rpb1* knock-out lines 25 dpi. Scale bar = 2 mm.
