## Supplemental Figure 8 for "Natural variation in Arabidopsis responses to *Plasmodiophora brassicae* reveals an essential role for RPB1"

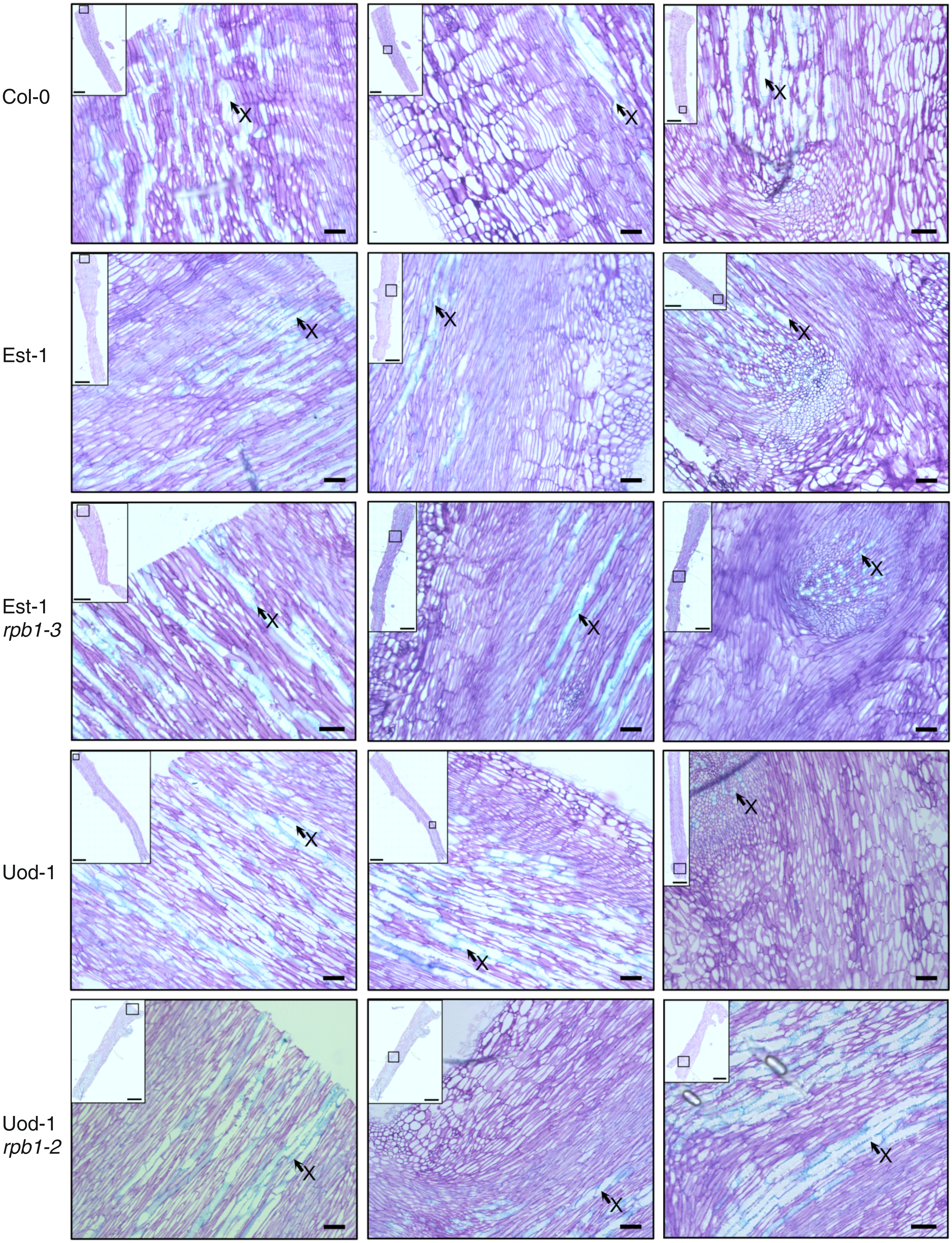


**Figure S8. Deletion of *RPB1* does not affect growth or development**.
Histological characterization of the hypocotyls of mock inoculated wildtype accessions and *rpb1* mutants 25 dpi for comparison with Figure 4. Micrographs in the left column focus on the uppermost central part of the hypocotyl, the central column focusses on the epidermal cells. In the right column the region of the interphase between the hypocotyl and the root is presented. X: Xylem cells. The scale bar in the inset overview picture corresponds to 1000 µm, the scale bar in the detailed pictures corresponds to 50 µm.
