## Supplemental Figure 9 for "Natural variation in Arabidopsis responses to *Plasmodiophora brassicae* reveals an essential role for RPB1"

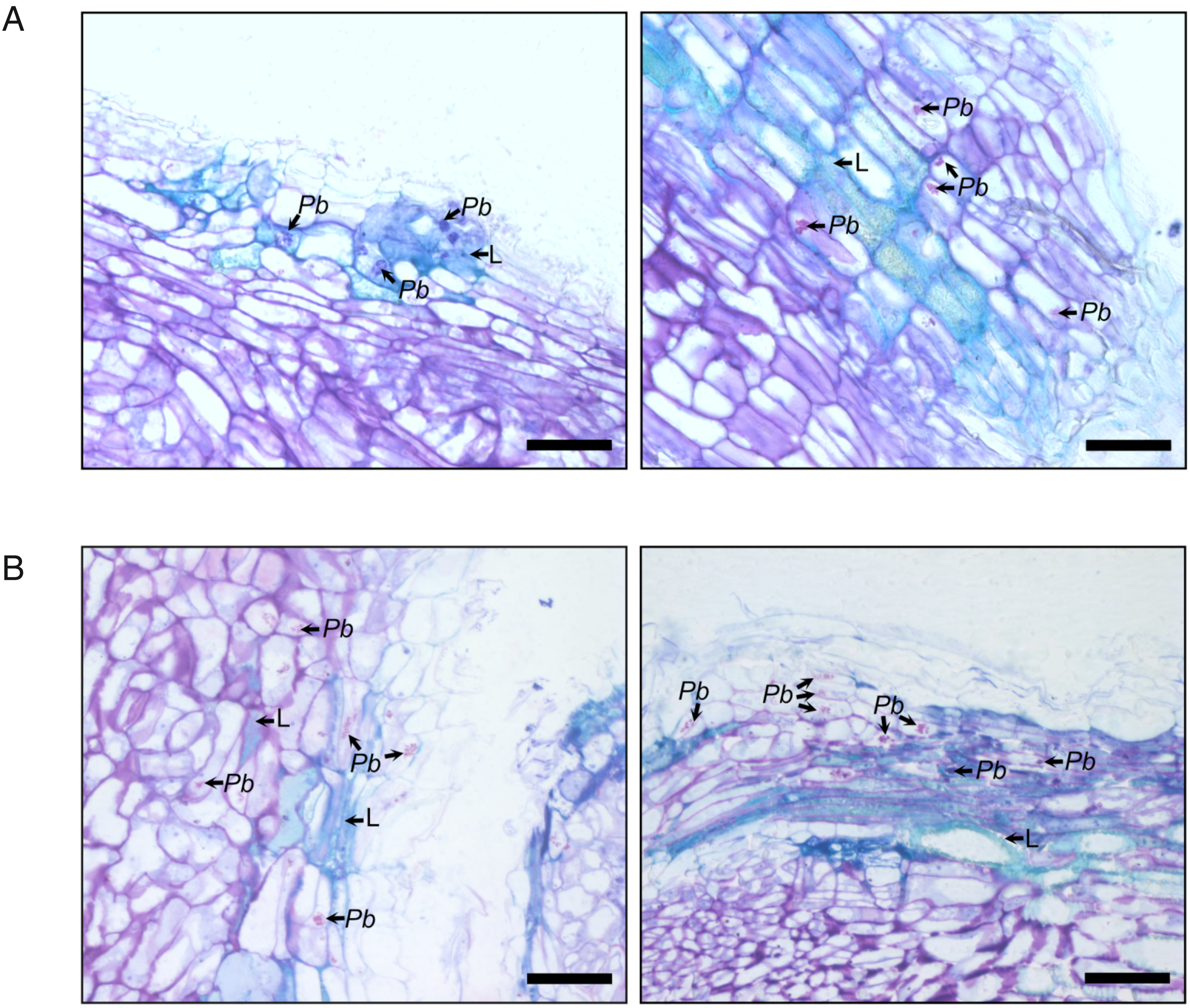


**Figure S9. *RPB1* is required for lignification of host cell walls in response to *P. brassicae*.**
Detailed visualisation of pathogen structures surrounded by lignified cells in the (A) Uod-1 and (B) Est-1 wildtype genotypes inoculated with *P. brassicae* 25 dpi. L: Lignified tissue, Pb: *P. brassicae* cells. The scale bar corresponds to 50 µm.
