## Supplemental Figure 10 for "Natural variation in Arabidopsis responses to *Plasmodiophora brassicae* reveals an essential role for RPB1"

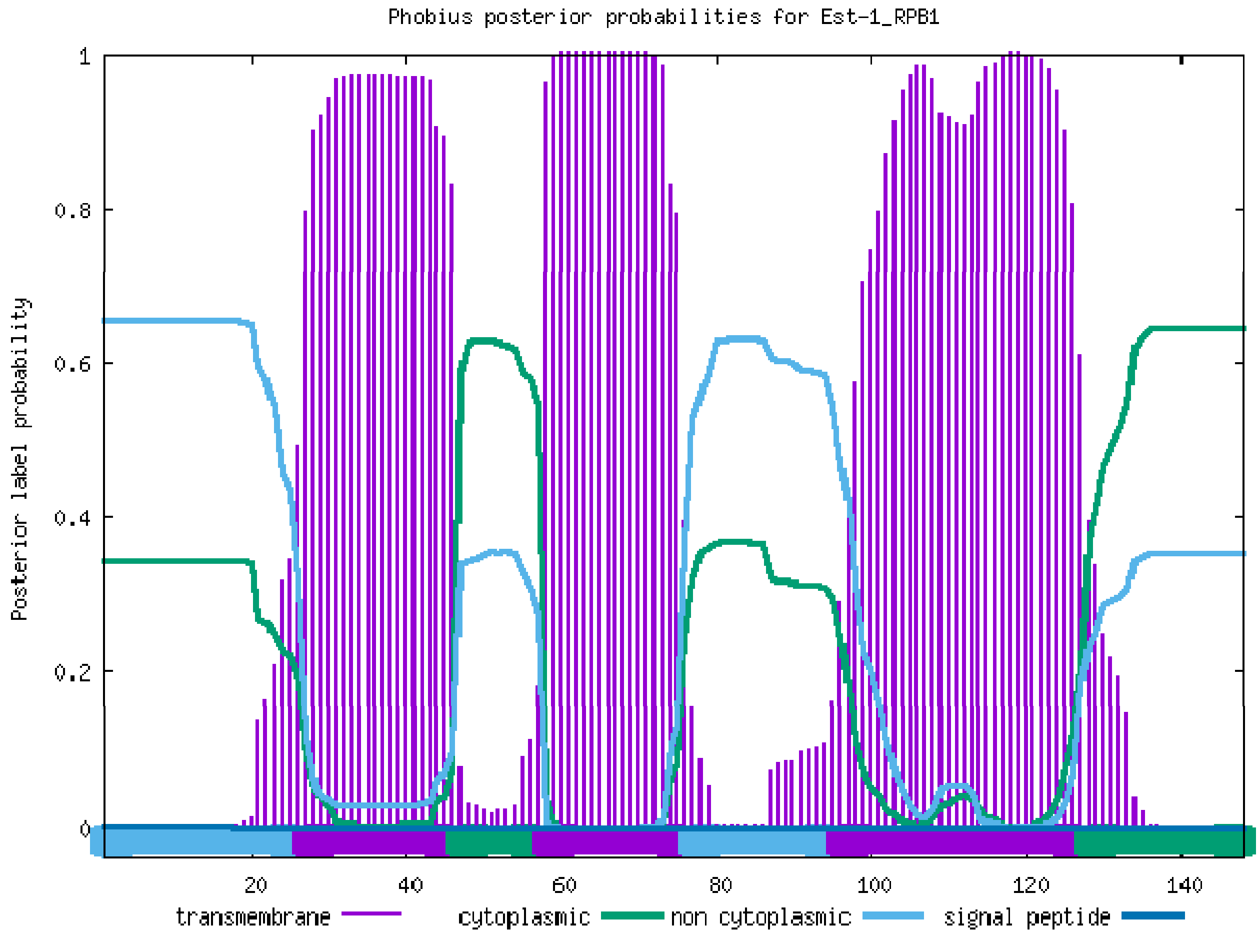


**Figure S10. RPB1 is predicted to contain membrane spanning domains.**
Prediction of membrane spanning domains (purple) in the sequence of RPB1 made by Phobius.
