## Supplemental Figure 11 for "Natural variation in Arabidopsis responses to *Plasmodiophora brassicae* reveals an essential role for RPB1"

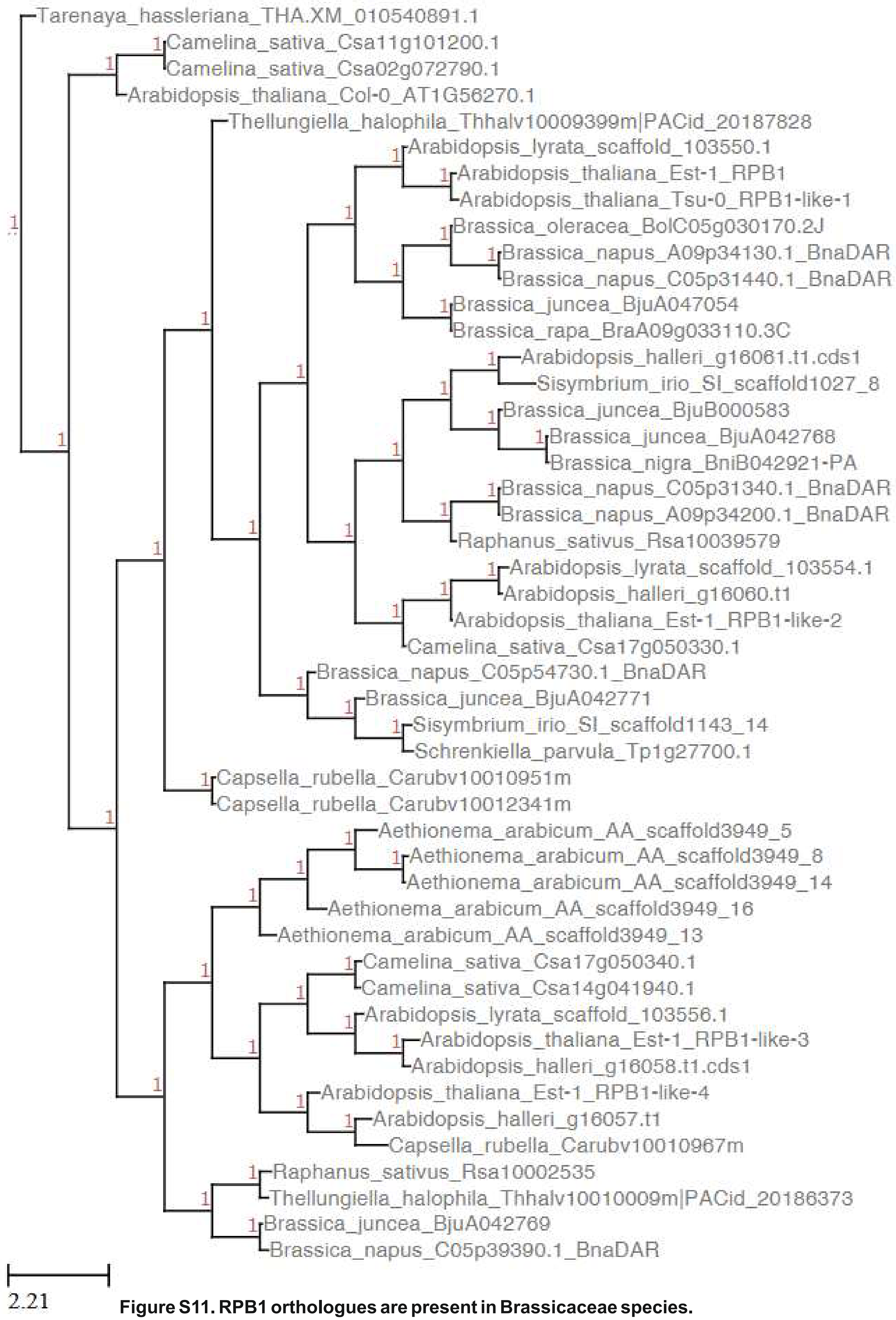


**Figure S11. RPB1 orthologues are present in Brassicaceae species.**
Identification of putative orthologues of *RPB1* and *RPB1-like* genes in Brassicacea species by OrthoFinder. All Arabidopsis *RPB1* related sequences were grouped in one orthogroup.
